# Centriole elimination during *C. elegans* oogenesis initiates with loss of the central tube protein SAS-1

**DOI:** 10.1101/2023.06.19.545600

**Authors:** Marie Pierron, Alexander Woglar, Coralie Busso, Keshav Jha, Tamara Mikeladze-Dvali, Marie Croisier, Pierre Gönczy

## Abstract

Centrioles are lost during oogenesis in most metazoans, ensuring that the zygote is endowed with the correct number of two centrioles, which are paternally contributed. How centriole architecture is dismantled during oogenesis is not understood. Here, we analyze with unprecedent detail the ultrastructural and molecular changes during oogenesis centriole elimination in *C. elegans*. Centriole elimination begins with loss of the so-called central tube and organelle widening, followed by microtubule disassembly. The resulting cluster of centriolar proteins then disappears gradually, usually moving in a microtubule- and dynein-dependent manner to the plasma membrane. Moreover, we find that neither Polo-like kinases nor the PCM, which modulate oogenesis centriole elimination in *Drosophila*, do so in *C. elegans*. Furthermore, we demonstrate that the central tube protein SAS-1 normally departs first from the organelle, which loses integrity earlier in *sas-1* mutants. Overall, our work provides novel mechanistic insights regarding the fundamental process of oogenesis centriole elimination.

## INTRODUCTION

Centrioles are small evolutionary conserved organelles characterized by a nine-fold radially symmetric arrangement of microtubules (reviewed in (Bornens, 2012; Gomes Pereira et al., 2021; Winey and O’Toole, 2014)). Centrioles play a crucial role as templates for the axoneme of cilia and flagella, which are critical for cell signaling and motility. A pair of centrioles surrounded by the pericentriolar material (PCM) constitutes the centrosome, the principal microtubule organizing center (MTOC) of animal cells. The number of centrioles is tightly regulated. Most proliferating cells are born with two centrioles, each of which then seeds the assembly of a procentriole during S-phase, leading to four centrioles during mitosis, two per spindle pole. Improper centriole number leads notably to defective spindle assembly and aberrant chromosome segregation, which can result in developmental failure and tumor progression.(reviewed in (Nigg and Holland, 2018)) How the rules governing centriole number control are tuned to serve specific physiological contexts is poorly understood.

Centriole number control must be modified at fertilization to ensure that the two gametes together contribute a total of two centrioles to the zygote. This requirement is achieved in most metazoan organisms by inactivating or eliminating maternally-derived centrioles and providing two centrioles from the sperm (reviewed in (Delattre and Gönczy, 2004; Manandhar et al., 2005)). There are two principal removal modes of maternally-derived centrioles. In the first mode, maternal centrioles are removed following the meiotic divisions (Crowder et al., 2015). This is the case in starfish, for example, where centrioles are present at spindle poles during the two female meiotic divisions, such that three maternal centrioles are eliminated through polar body extrusion, with the last one being removed in the zygote (Borrego-Pinto et al., 2016; Pierron et al., 2020; Shirato et al., 2006; Sluder et al., 1993a, 1993b). In the second and most prevalent mode, which occurs notably in worms, flies and vertebrates, maternal centrioles are removed during oogenesis, before the meiotic divisions. A mechanism underlying this removal mode has been uncovered in *Drosophila*, where loss of the Polo kinase from centrosomes during oogenesis results in PCM disassembly and subsequent loss of focused centriolar proteins (Pimenta-Marques et al., 2016). In *C. elegans*, depletion of the RNA helicase CGH-1 or the TRIM-NHL protein LIN-41 delays, but does not abolish, oogenesis centriole elimination, as deduced from the prolonged persistence of foci containing centriolar proteins (Matsuura et al., 2016; Mikeladze-Dvali et al., 2012). Importantly, the studies to date did not explain how these molecular players results in centriole disassembly, nor did they assess organelle integrity at the ultrastructural level. Analysis by electron-microscopy (EM) is critical to ensure that foci bearing centriolar proteins correspond to *bona fide* centrioles. This is exemplified by the situation in mice, where foci bearing the centriolar marker Centrin2::GFP are present at the poles of the female meiotic spindle until metaphase II, despite centrioles being absent as judged by EM (Simerly et al., 2018). Overall, despite being of fundamental importance for successful sexual reproduction, centriole alterations during oogenesis have not been analyzed in detail at the ultrastructural level, and understanding of the mechanisms governing this process is incomplete.

By contrast to the paucity of information regarding the sequence of events and the mechanisms underlying oogenesis centriole elimination, substantial knowledge has been accrued regarding the evolutionary conserved proteins that assemble the organelle (reviewed in (Banterle and Gönczy, 2017; Gomes Pereira et al., 2021; Nigg and Holland, 2018; Ohta et al., 2017)). In *C. elegans*, the first components recruited to the assembly site are the interacting proteins SAS-6 and SAS-5, which form a so-called inner tube, the equivalent of the cartwheel hub in other systems. SAS-6/SAS-5 recruitment requires the function of the ZYG-1 kinase, as well as the presence of SAS-7 and SPD-2 on the pre-existing centriole. Next comes SAS-4, which is thought to facilitate the processive elongation of the microtubule singlets that are then added to complete organelle assembly. The precise localization of 12 centriolar and PCM core components has been uncovered in meiosis prophase I using Ultrastructure-Expansion coupled to STED super-resolution microscopy (U-Ex-STED), enabling also the mapping of these proteins onto specific ultrastructural elements defined by EM (Fig. 1A) (Woglar et al., 2022). This work established notably that the microtubule interacting protein SAS-1 localizes to the so-called central tube, a circular element between the SAS-6/SAS-5-containing inner tube and microtubules. *sas-1* is dispensable for centriole assembly, but essential to maintain stability of sperm-contributed centrioles in the zygote and of centrioles during subsequent embryogenesis (von Tobel et al., 2014). Furthermore, the U-Ex-STED map showed that the centriolar proteins HYLS-1, SPD-2, as well as the PCM core components PCMD-1 and SPD-5, all localize peripheral to the microtubules, onto the so-called paddlewheels (Woglar et al., 2022). The centriolar protein SAS-7, as well as the PCM component γ-tubulin (TBG-1), localize more externally still. TBG-1, together with its interactors GIP-1 and GIP-2, as well as its partner MZT-1, is present in the PCM core during interphase, with more protein being present during mitosis, thereby enabling robust microtubule nucleation. Other proteins, including the ZYG-9/TAC-1 complex, PLK-1 and the Aurora A kinase AIR-1, contribute to centrosome maturation and microtubule nucleation during mitosis (reviewed in (Pintard and Bowerman, 2019)).

**Figure 1.**
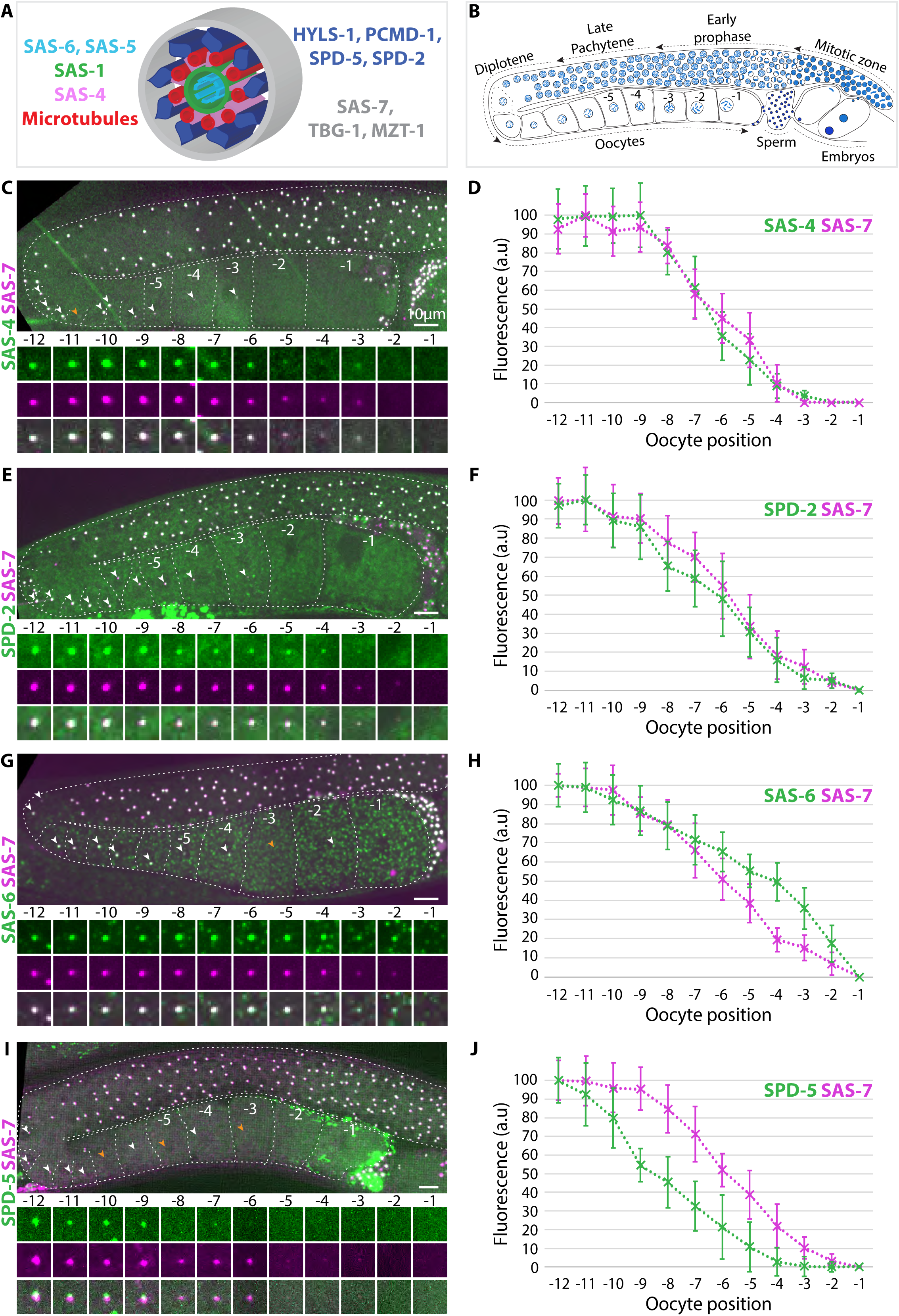
Progressive loss of foci enriched in centriolar and PCM components. **A.** Schematic of a slightly tilted view of worm centriole, illustrating the distribution of select centriolar and PCM components. The dimensions of meiotic prophase centrioles are ∼140nm in diameter x ∼100 nm in height. Modified after (Woglar et al., 2022). **B.** Schematic of an adult hermaphrodite gonad showing progression of germ cells from the mitotic zone, through early meiotic prophase, late pachytene and then diplotene, before cellularization into maturing oocytes. Oocytes are numbered in reverse order according to their distance from the spermatheca. The most mature −1 oocyte will enter the spermatheca where it is fertilized, thereby giving rise to the zygote. The condensation state of the chromatin is schematized in blue. Not drawn to scale. **C-J.** Gonads from synchronized worm populations expressing the indicated two markers (C, E, G, I), together with corresponding fluorescence intensity quantifications (D, F, H, J, in arbitrary units normalized to the maximal and minimal fluorescence intensity in analyzed oocytes in each case ± SD). Here and in Fig.3, we measured the correlation for each pair of markers and R^2^ of fits are indicated together with N for each panel. (C, D): SAS-4::mCherry and GFP::SAS-7, N=10 and R^2^=0.98; (E, F): SPD-2::GFP and RFP::SAS-7, N=9 and R^2^=0.98; (G, H): SAS-6::GFP and RFP::SAS-7, N=10 and R^2^=0.93; (I, J) RFP::SPD-5 and GFP::SAS-7, N=12 and R^2^=0.84. Unless stated otherwise, here and in all figures, images are max-intensity projections of relevant z-slices from 3D confocal live imaging. Dashed white lines delineate the gonad and oocytes, which are numbered as in B. Arrowheads point to the location of centriolar protein foci. Here, as well as in Fig. 2 and Fig. 3, orange arrowheads point to foci that are not obvious in the overview because they reside in deeper slices. Insets are 12 microns thick max-intensity projections. Scale bars are indicated in a single image per Figure if all images are scaled identically.

*C. elegans* oocytes are produced in an assembly line fashion that provides access to all stages of the centriole elimination process within a single gonad in a spatial-temporal gradient (Fig. 1B)(Lints and Hall, 2004; Pazdernik and Schedl, 2013). Germ cell nuclei are produced in the mitotic zone located in the distal end of the gonad. Nuclei then progress towards the proximal end, going through early prophase, late pachytene and then diplotene of meiosis I, before undergoing cellularization after the loop region, yielding oocytes that then mature further. Fertilization occurs as fully mature oocytes pass through the spermatheca and execute the two meiotic divisions without maternal centrioles. Prior analysis of *C. elegans* oogenesis centriole elimination using antibodies against centriolar and PCM proteins, as well as transgenic worms expressing fusion proteins, combined with serial-section EM of a few cells, uncovered that centrioles are eliminated during diplotene (Mikeladze-Dvali et al., 2012). However, the sequence of ultrastructural changes during centriole elimination has not been addressed in this initial study. Moreover, the prior work was conducted before the advent of expansion microscopy, which holds the potential to provide high spatial resolution information on changes in molecular architecture during this process.

Here, we set out to investigate in more depth the mechanisms underlying centriole elimination during *C. elegans* oogenesis. We conducted live imaging of endogenously tagged centriolar and PCM core proteins, including in a microfluidic device enabling long term monitoring. We uncovered that the signal intensity of centriolar protein foci starts to diminish in diplotene, but remains detectable in maturing oocytes, where it usually moves to the plasma membrane. Moreover, we established that neither the Polo like kinases PLK-1, PLK-2 and PLK-3, nor the PCM, protects centrioles from elimination in *C. elegans* oogenesis. Furthermore, combining Correlative Light Electron Microscopy (CLEM) with U-Ex-STED, we uncovered discrete changes in centriole architecture during centriole elimination, and correlated them with corresponding alterations in protein abundance. In particular, we found that the initial step of elimination is marked by the departure of SAS-1 from the centriole and an increase of organelle diameter. Moreover, we show that SAS-1 plays a critical role in timing centriole elimination during worm oogenesis.

## RESULTS

### Progressive loss of centriolar and PCM protein foci

We set out to analyze the process of centriole elimination during *C. elegans* oogenesis in new detail. First, we investigated whether centriolar and PCM proteins are removed at the same time or instead in a sequence that may be indicative of the underlying mechanism. To this end, we utilized worm strains simultaneously expressing fluorescently tagged endogenous SAS-7 and a tagged version of another centriolar or PCM protein expressed from the endogenous locus or as a validated transgene (Table 1). We performed live confocal imaging of young adult hermaphrodites, quantifying fluorescence intensities in foci enriched in centriolar or PCM proteins from the diplotene stage in the loop region until the last oocyte before the spermatheca (Fig. 1; Fig. S1). Note that by convention oocytes are numbered according to their position relative to the spermatheca, with the most proximal one referred to as the −1 oocyte (Fig. 1B). We found that the fluorescence intensity of foci enriched in centriolar or PCM proteins associated with each germ cell nucleus starts to decay in diplotene, typically in the −10 oocyte (Fig. 1 D, F, H, J). Moreover, we found dimmer foci as far as the −2 oocyte for the centriolar proteins SAS-6, SAS-4, SAS-7 and HYLS-1 (Fig. 1C, 1G; Fig. S1A). The same was true for the PCM core components SPD-2, SPD-5 and PCMD-1, as well as the γ-tubulin-interacting proteins GIP-1 and MZT-1 (Fig. 1E, 1I; Fig. S1B-D). Overall, we conclude that all centriolar and PCM core proteins analyzed diminish in intensity starting approximately in the −10 oocyte, but remain detectable in a focus until later.

**Table 1:**
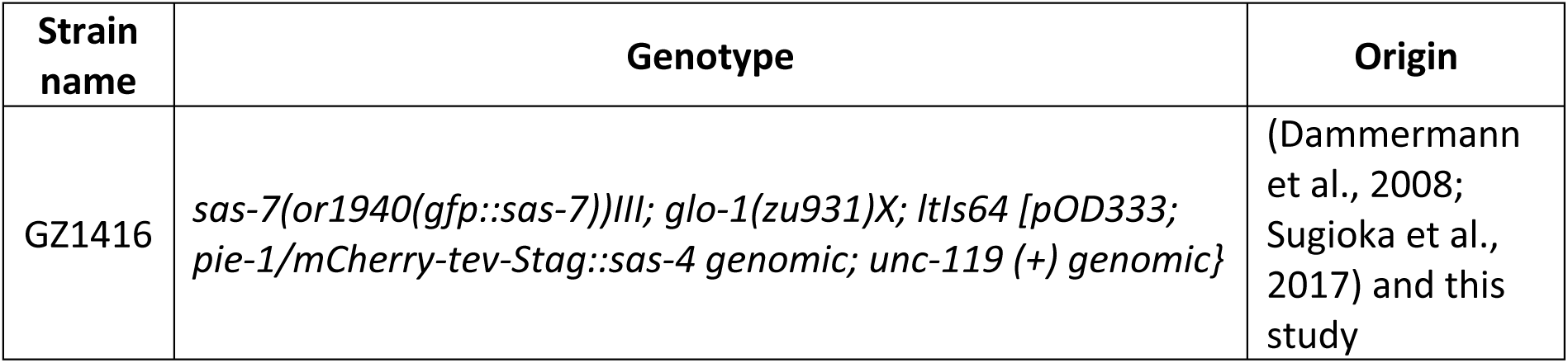
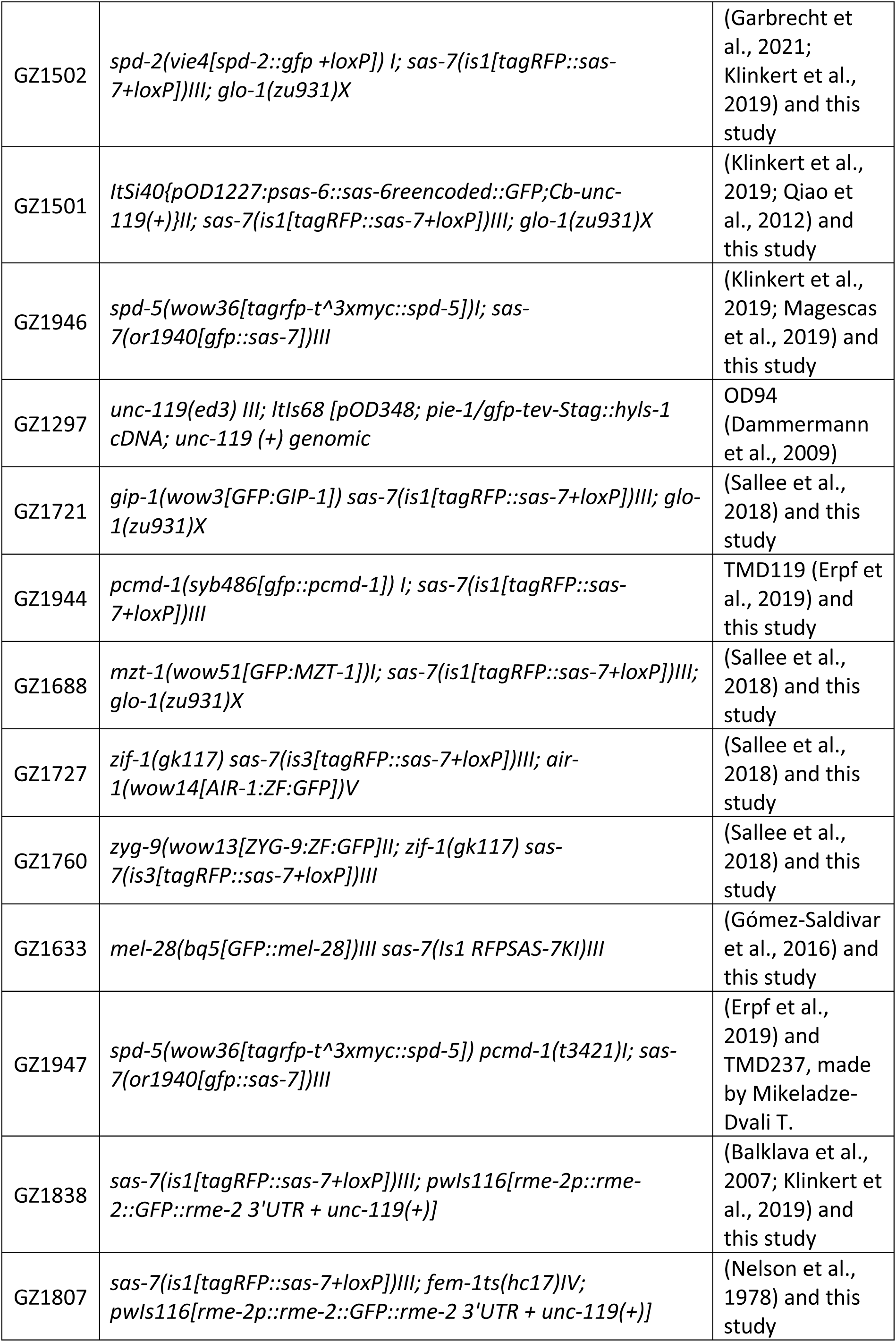
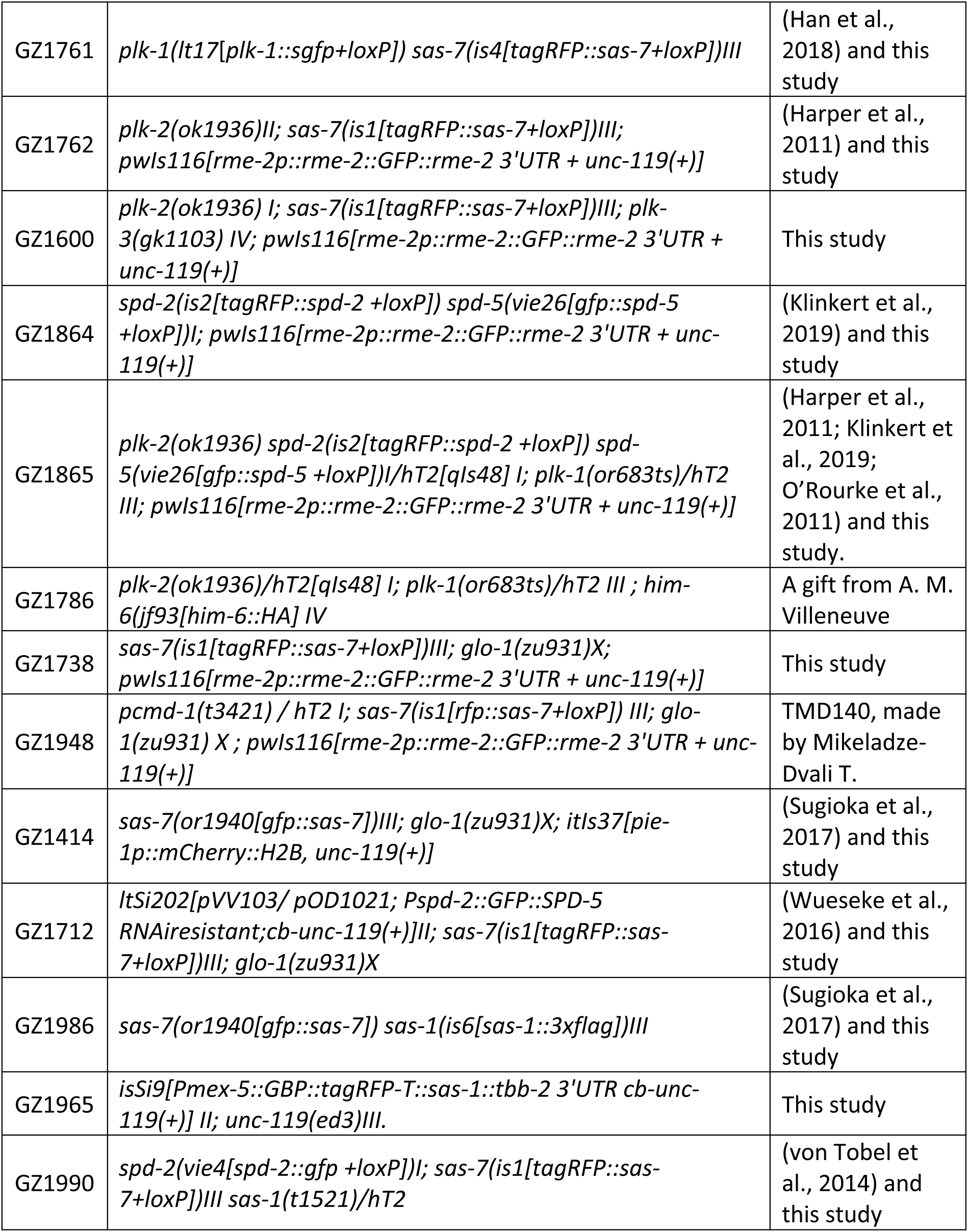
list of strains used in this study.

We next analyzed fluorescence intensity decay of proteins in the focus with respect to SAS-7, which served as a standard in all animals. With the exception of SPD-5 and SAS-6, we found that focused fluorescence intensity decreases similarly for SAS-7 and the other proteins analyzed, indicating concomitant decay (Fig. 1D, 1F; Fig. S1A-S1D). In the case of SAS-6, the cytoplasm of −6 to −1 oocytes contains many dim foci (Fig. 1G), which prevents one from thoroughly assessing whether a focus of centriolar SAS-6 persists after disappearance of the SAS-7 focus, although slower decay of SAS-6 fluorescence levels suggests that this might be the case (Fig. 1H). By contrast, we found that the PCM core protein SPD-5 is removed earlier than SAS-7 (Fig. 1I, 1J). Furthermore, MZT-1 and GIP-1 remain present in a focus in maturing oocytes despite the absence of SPD-5 (Fig. S1C, S1D), which is different from the situation in embryos, where SPD-5 is required for γ-tubulin recruitment to centrosomes (Hamill et al., 2002). We also found that centrosomal AIR-1 and ZYG-9 are present only in mitotic cells in the distal region of the gonad, but not thereafter (Fig. S1E, S1F), as anticipated from their recruitment to mature centrosomes in the early embryo (Matthews et al., 1998; Schumacher et al., 1998), as well as from analogous distributions in other systems (Crosio et al., 2002; Giet et al., 2002).

Together, these data reveal that the removal of centriolar and PCM proteins occurs progressively during oogenesis, starting in diplotene of meiotic prophase I and finishing in mature oocytes. Most proteins depart in a concomitant manner, except for SPD-5, which is removed earlier, and SAS-6, which appears to be removed later.

### Foci containing centriolar proteins detach from the nucleus and move to the plasma membrane in a microtubule- and dynein-dependent manner

Whilst analyzing the distribution of centriolar and PCM core protein foci, we noticed that instead of residing in the vicinity of the nucleus, as is usually the case for centrioles, such foci are sometimes present in the cytoplasm or at the plasma membrane, in particular in the last few oocytes (see Fig. S1A, −5 oocyte, and Fig. S1B, −3 oocyte). To investigate this phenomenon further, we utilized a microfluidic device designed for long-term non-invasive imaging of *C. elegans*, allowing us to track the focus of centriolar proteins for hours (Berger et al., 2018). We performed 4D widefield live imaging of gonads expressing RFP::SAS-7 and MEL-28::GFP, a nucleoporin localized at the nuclear envelope (Galy et al., 2006), or else expressing RFP::SAS-7 and RME-2::GFP, a yolk receptor enriched at the plasma membrane of the most mature oocytes (Grant and Hirsh, 1999)(Fig. 2A; see Materials and methods). We imaged 11 gonads, starting with oocytes situated in positions −6 to −4 at the onset of the experiment, following them for 3-5 hours during their progression towards the spermatheca. This analysis established that in ∼80% of cases (26/33 oocytes), the RFP::SAS-7 focus detaches from the nuclear envelope (Fig. 2A, 0’), and moves towards the plasma membrane (Fig. 2A, 52’). The focus persists in that location during oocyte enlargement (Fig. 2A, 156’), before becoming undetectable (Fig. 2A, 166’). We generated kymographs from 7 movies in which the focus of RFP::SAS-7 remained largely in the XY imaging axis, finding an average velocity of ∼0.65μm/min (Fig. 2B, 2C) (see Discussion).

**Figure 2.**
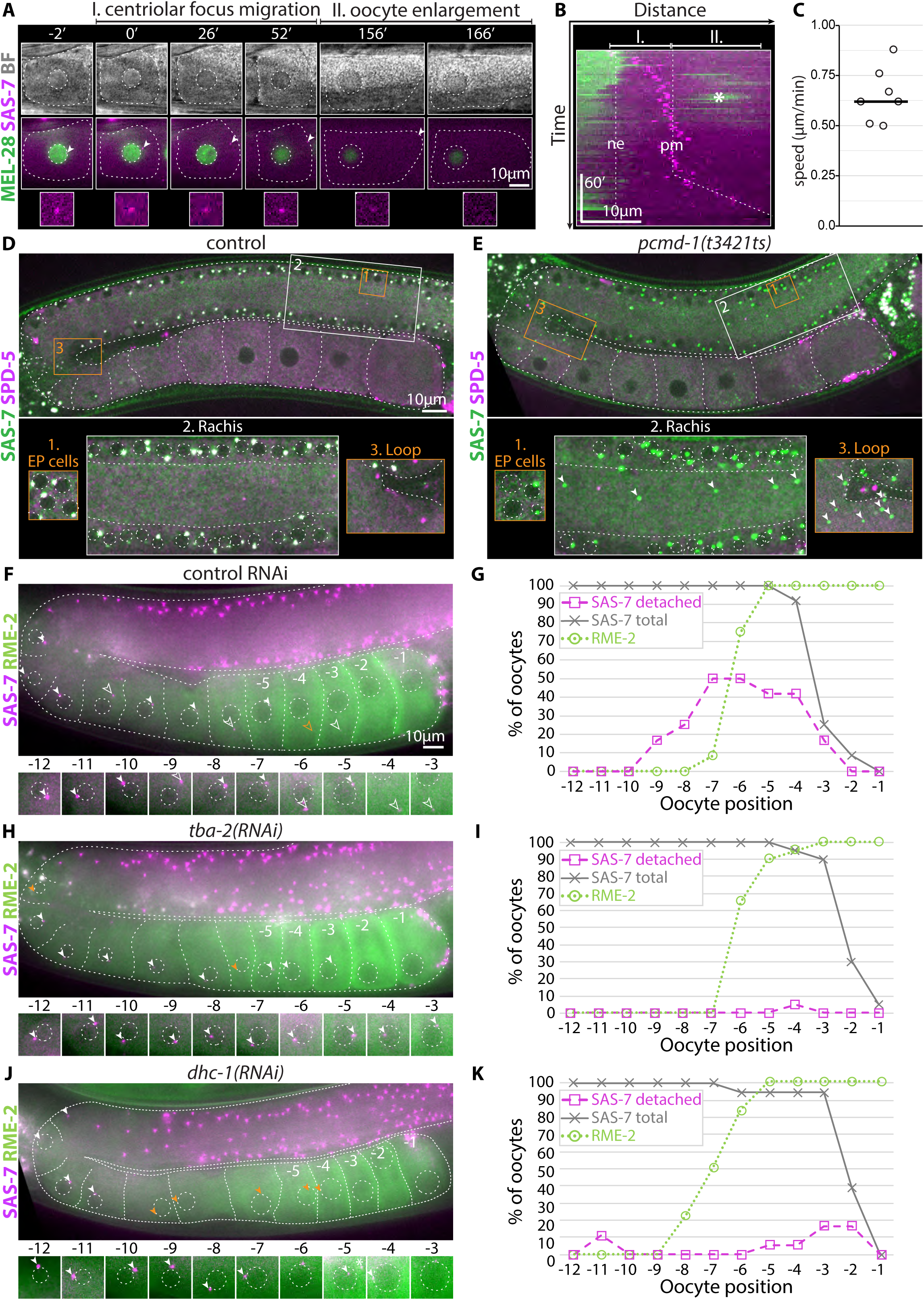
Centriolar foci remnant detach from the nuclear envelope and move to the plasma membrane in a microtubule- and dynein-dependent manner. **A.** Widefield time lapse imaging of an oocyte initially located at position −6, which reaches position −1 position by the end of the movie, from a worm immobilized in the microfluidic device expressing MEL-28::GFP and RFP::SAS-7. Top: brightfield (BF); middle: merge between MEL-28::GFP (green) and RFP::SAS-7 (magenta); bottom: magnified insets showing RFP::SAS-7 focus indicated by an arrowhead at each time point. Brightfield images and MEL-28::GFP signal were used to outline the plasma membrane and the nuclear envelope, respectively, both marked with dashed lines. Note RFP::SAS-7 centriolar focus detaching from the nuclear envelope (0’), moving to the plasma membrane (0’-52’), and persisting there during oocyte enlargement (156’), before disappearing (166’). Time in minutes. See also Movie S1. **B.** Kymograph from widefield time-lapse imaging in A and Movie S1, showing movement of the RFP::SAS-7 focus (magenta) away from the nuclear envelope (ne) towards the plasma membrane (pm) (I.). Note that the RFP::SAS-7 focus remains at the plasma membrane and is gradually more distant from the nucleus due to oocyte enlargement (II.), before disappearance. MEL-28::GFP signal is dimmer at times because the nucleus moves deeper in the tissue (see also Movie S1). Asterisk marks the nucleus of the adjacent oocyte present transiently in the vicinity. **C.** Quantifications of RFP::SAS-7 focus movement velocity. Oocytes were between the −4 and the −6 position at the beginning of the movies and imaged at least until the RFP::SAS-7 focus reached the plasma membrane. Measurements were performed when the RFP::SAS-7 focus moved mainly within the xy plane to ease quantification. Horizontal line: average speed. N= 7 oocytes. **D, E.** Gonads from control worms (D) and *pcmd-1*(*t3412ts*) mutant animals at the restrictive temperature (E), both expressing GFP::SAS-7 and RFP::SPD-5. Highlighted regions on the top (1: early pachytene, EP; 2: rachis; 3: loop region) are shown magnified at the bottom as max intensity projections of relevant z-slices, which can be different from those in the overview image. Note absence of SPD-5 in *pcmd-1*(*t3412ts*) mutant animals as evidenced by the presence of green foci. Note also that some nuclei lack centrioles already in early pachytene, indicating that centrioles detach precociously, accumulating in the rachis and the loop region. In the insets, dashed lines delineate the rachis and loop borders, dashed circles nuclei, whereas arrowheads point to mislocalized centriolar foci. **F-K.** Widefield images of gonads from worms of indicated RNAi conditions expressing RFP::SAS-7 and RME-2::GFP (F, H, J) and corresponding quantifications (G, I, K). The presence of centriolar foci was quantified (G, I, K: grey lines -SAS-7 total), distinguishing those at the nuclear envelope (F, H, J: arrowheads) or detached from it (F,H,J: open arrowheads and G, I, K: magenta dashed lines -SAS-7 detached). RME-2::GFP becomes progressively enriched at the plasma membrane as the oocyte matures (see Fig.3B for confocal images). Note that disappearance of centriolar foci occurs in a timely manner in all conditions as shown by SAS-7 focus disappearance relative to oocyte maturation as quantified with RME-2::GFP increase. In J, −5 oocyte, asterisk marks RFP::SAS-7 from a neighboring somatic cell. N: G=12; I=18; K=20.

What could cause movement of the RFP::SAS-7 focus towards the plasma membrane? Protein aggregates are known to move within mature oocytes before being degraded via a sperm-derived signal (Adam Bohnert and Kenyon, 2017). However, we found that RFP::SAS-7 foci movement was not altered in feminized mutant animals lacking sperm (Fig. S2A-S2D). Moreover, we investigated whether the PCM could facilitate juxtanuclear retention of RFP::SAS-7 foci. Compatible with this notion, centriolar foci detach prematurely from germ cell nuclei in *pcmd-1(t3421ts)* mutant animals, which fail to assemble PCM (Erpf et al., 2019); as a result, centriolar foci devoid of nuclei are present in the gonad rachis and the loop region (Fig. 2D, 2E). This finding suggests that the PCM contributes to linking centrioles to the nuclear envelope. In turn, this may explain why RFP::SAS-7 foci normally depart from the nuclear envelop after the loop region, where SPD-5 levels decrease substantially (see Fig. 1I, 1J).

Next, we used RNAi in worms expressing RFP::SAS-7 and RME-2::GFP to test whether microtubule and F-actin, as well as molecular motors, are required for focus movement towards the plasma membrane. To increase throughput, we imaged worms at a single time point, assessing whether centriolar foci were juxtanuclear or not in −12 to −1 oocytes (Fig. 2F-2K; Fig. S2E-S2I). In line with the live imaging observations, we found that centriolar foci were present in the cytoplasm in approximately 50% of −7 oocytes in the control condition (Fig. 2F, 2G). No change was observed either upon F-actin depletion, despite modifications of gonad architecture following *act-1*(*RNAi*) (Fig. S2E, S2F). In stark contrast, depletion of microtubules through *tba-2*(*RNAi*), as well as of the dynein heavy chain DHC-1 or the dynein light chain DLC-1, essentially abrogated movement, with RFP::SAS-7 foci almost invariably remaining at the nuclear envelope (Fig. 2H-2K; Fig. S2G). Depletion of the conventional kinesin UNC-116 led to a milder reduction (Fig. S2H, S2I). Therefore, microtubules and dynein are required for RFP::SAS-7 focus movement to the plasma membrane, with kinesin contributing as well.

Overall, our findings uncover that oogenesis centriole elimination in *C. elegans* comprises a late phase during which the focus containing diminishing levels of centriolar and PCM proteins can detach from the nucleus and move towards the plasma membrane in a microtubule- and dynein-dependent manner. Importantly, however, in approximately 20% of oocytes, the focus of RFP::SAS-7 becomes undetectable in the vicinity of the nucleus, without apparent movement towards the plasma membrane. Moreover, when microtubules or dynein components were depleted, RFP::SAS-7 centriolar foci eventually disappeared (see Fig. 2I, 2K, grey lines), despite the absence of plasma membrane-directed movement. Therefore, even if frequent in the control condition, this phenomenon is not essential for the clearance of foci with centriolar and PCM proteins during *C. elegans* oogenesis.

### Depletion of Polo-like kinases or PCM removal does not lead to precocious centriole elimination during *C. elegans* oogenesis

Given that Polo-mediated PCM removal triggers oogenesis centriole elimination in the fly (Pimenta-Marques et al., 2016), we investigated whether Polo-like kinases and the PCM operate similarly in the worm. There are three Polo-like kinases in *C. elegans*: PLK-1, which is essential and closest to Polo, exerting analogous functions during centrosome maturation and embryonic cell division (Chase et al., 2000); PLK-2, which is not essential and functions in meiotic chromosome organization (Brandt and Kim, 2021); the more divergent and non-essential PLK-3, which has no ascribed function. Previous immunostaining established that PLK-1 and PLK-2 in the gonad localize at centrosomes exclusively in the mitotic zone (Harper et al., 2011). Accordingly, we found that endogenously tagged PLK-1::GFP marks centrosomes exclusively in dividing cells in that region (Fig. 3A).

**Figure 3.**
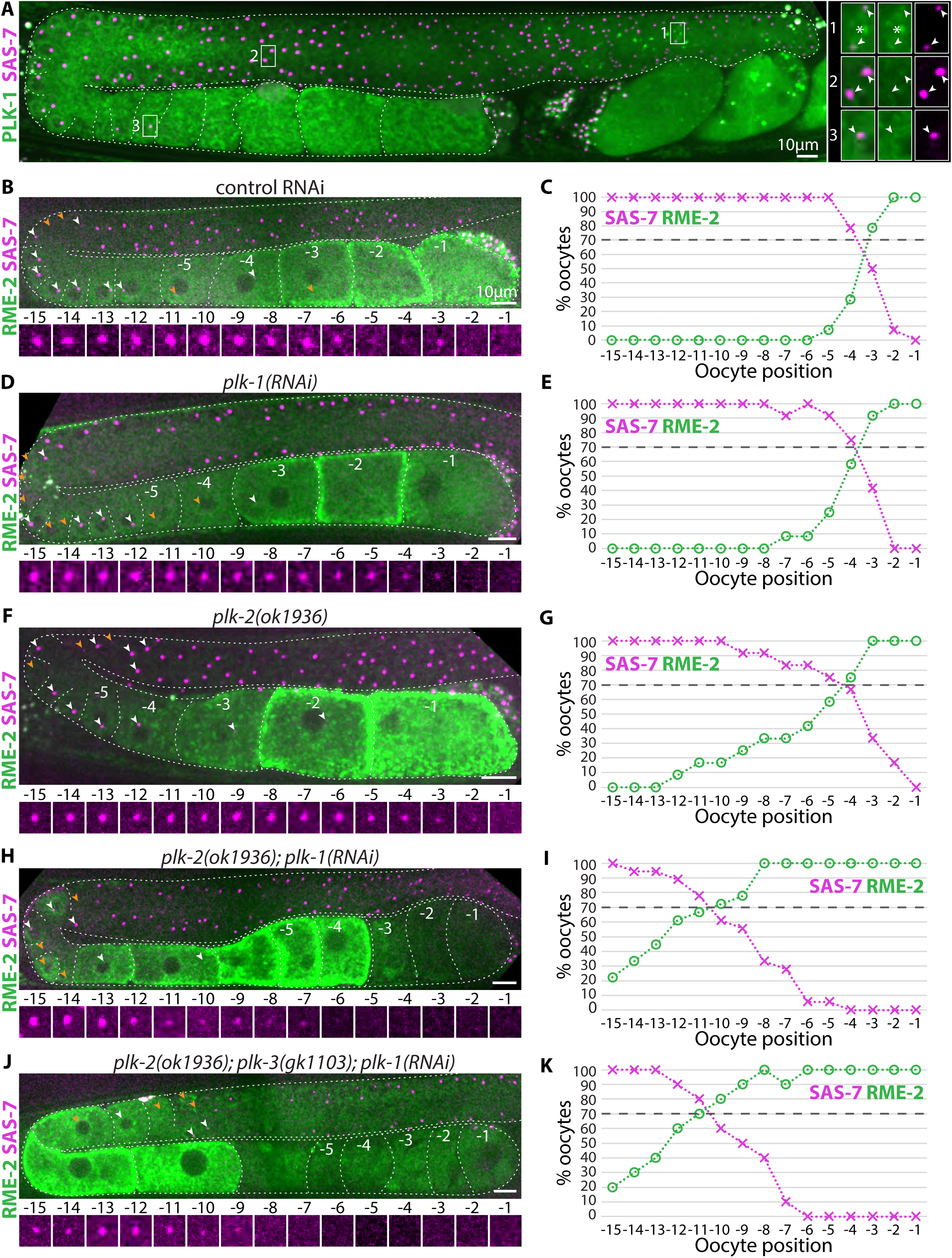
Depletion of PLKs does not lead to precocious centriole elimination. **A.** Gonad expressing PLK-1::GFP and RFP::SAS-7. White squares highlight three regions magnified as insets on the right. Arrowheads point to SAS-7 foci (magenta) and their corresponding position in the PLK-1 channel (green). Asterisk marks PLK-1 on chromosomes. Note that centriolar PLK-1 is present exclusively during mitosis (inset 1). **B-K.** Gonads from worms of indicated genotypes and RNAi conditions expressing RME-2::GFP and RFP::SAS-7 (B, D, F, H, J), together with corresponding quantifications (C, E, G, I, K). Note that in the double and triple mutant/RNAi conditions, RFP::SAS-7 decay and RME-2::GFP enrichment at oocytes plasma membrane are shifted toward the loop but still occur in a concomitant manner, approximately when 70% of oocytes contain both markers (horizontal dashed line in the panels on the right). C : N=14 and R^2^=0.96; E : N=12 and R^2^=0.91; G : N=12 and R^2^=0.87; I :N=18 and R^2^=0.86; K=10 and R^2^=0.81.

In principle, PLK-1 could be present below detection levels in later stages when elimination occurs, or exerts a function in the mitotic zone that translates into subsequent organelle elimination. To explore these formal possibilities, we set out to investigate the role of PLK-1, as well as PLK-2 and PLK-3 in case redundancies were at play. We used *plk-1(RNAi)* feeding conditions that result in meiotic arrest (Budirahardja and Gönczy, 2008), as well as the null alleles *plk-2*(*ok1936*) and *plk-3*(*gk1103*). All analyzed animals expressed RFP::SAS-7 and RME-2::GFP to score centriolar focus disappearance relative to oocyte maturation, using the presence of RME-2::GFP on the plasma membrane to this end (Materials and methods). Importantly, we found that the timing of RFP::SAS-7 centriolar focus elimination is comparable in *plk-1(RNAi)* animals and control worms, with ∼70% of −3/-4 oocytes harboring both RFP::SAS-7 foci and RME-2::GFP (Fig. 3B-3E). In *plk-2*(*ok1936*) mutant gonads, oocyte maturation occurs earlier than in the control, as evidenced by the presence of RME-2::GFP positive oocyte as early as positions −12/-11 (Fig. 3F, 3G). RFP::SAS-7 foci are also sometimes lost earlier, but the concordance between RME-2::GFP rise and RFP::SAS-7 decay is preserved, demonstrating that the timing of elimination is not altered (Fig. 3G). When PLK-1 is depleted by RNAi in *plk-2*(*ok1936*) animals, oocyte maturation is even more precocious, but RFP::SAS-7 disappearance still occurs after RME-2::GFP increase (Fig. 3H, 3I). Finally, *plk-1(RNAi) plk-2*(*ok1936*) *plk-3*(*gk1103*) triply inactivated animals do not exhibit additional phenotypes regarding oocyte maturation or centriole elimination timing (Fig. 3J, 3K).

Because RNAi-mediated depletion can sometimes be incomplete, we combined *plk-2(ok1936)* with the thermosensitive allele *plk-1(or683ts)* at the restrictive temperature to achieve the strongest possible depletion condition compatible with life. This yielded highly disorganized gonads, yet with seemingly normal centriole elimination timing, as SPD-2 and SPD-5 foci persist until the RME-2::GFP signal appears, as in control worms (Fig. S3A, S3B). In order to further test whether centriole elimination in *plk-2(ok1936) plk-1(or683ts)* animals correlates with oocyte maturation, we immunostained gonads with IFA-1 pan-centriolar antibodies and counterstained them with a DNA dye to assess chromosome condensation as a proxy for meiotic progression (see Fig. 1B) (Phillips et al., 2009). We found in both control and mutant worms that centriolar foci are present in the vicinity of nuclei with chromosomes characteristic of the late diplotene stage (Fig. S3C, S3D). Overall, we conclude that Polo-like kinases do not modulate centriole elimination timing in *C. elegans*.

We next tested whether the PCM may be required for centriole stability independently of PLK-1, PLK-2 and PLK-3. PCMD-1 is essential for generating the PCM core (Erpf et al., 2019), and we therefore utilized the strong *pcmd-1*(*t3421ts*) mutant allele at the restrictive temperature to test whether PCM removal results in premature loss of centriolar foci. Importantly, we found that although RFP::SPD-5 was absent from centrosomal foci during oogenesis in *pcmd-1*(*t3421ts*) animals, as anticipated from findings in the embryo (Erpf et al., 2019), the kinetics of GFP::SAS-7 or SAS-4 loss were not altered (Fig. 4A-C; Fig. S3E, S3F). Moreover, we found that GFP::SAS-7 decay in such animals coincides with RME-2::GFP enrichment, further demonstrating that elimination timing is not impacted in the absence of SPD-5 (Fig. 4D). We conclude that PCM loss does not lead to precocious loss of foci with centriolar and PCM core proteins during *C. elegans* oogenesis.

**Figure 4.**
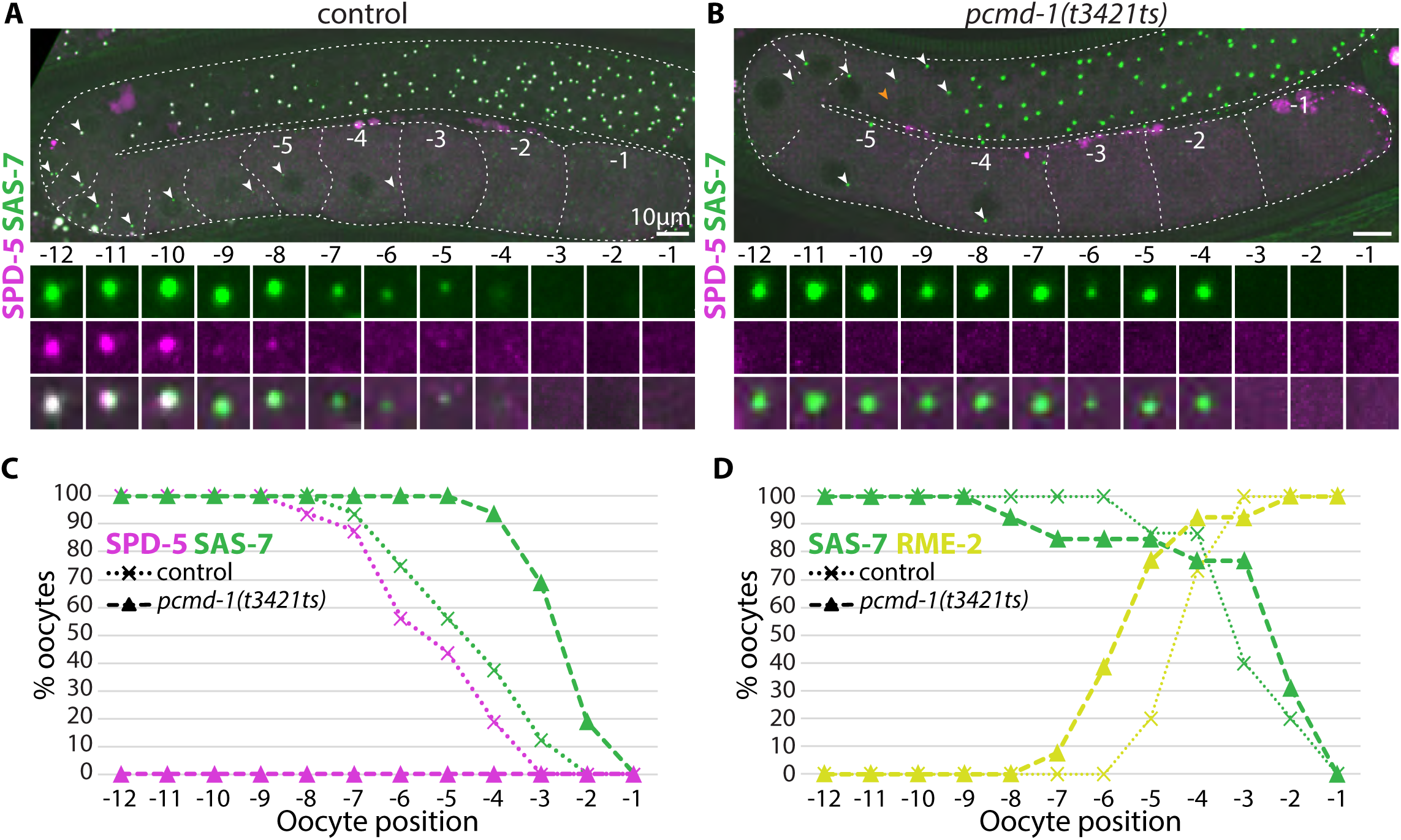
Removal of the PCM core component SPD-5 does not impact centriole elimination timing. **A-C.** Gonads from worms of indicated genotypes expressing RFP::SPD-5 and GFP::SAS-7 (A, B) and corresponding quantifications (C), illustrating persistence of SAS-7 foci despite absence of SPD-5 in the *pcmd-1*(*t3421ts*) mutant at the restrictive temperature. Control: N=16; *pcmd-1(t3421ts)*: N= 16. **D.** Percentage of oocytes bearing SAS-7 foci and RME-2::GFP enriched at the plasma membrane at indicated positions in control and *pcmd-1*(*t3421ts*) mutant worms at the restrictive temperature. Control: N=15; *pcmd-1(t3421ts)*: N=13.

### Central tube loss and centriole widening mark the onset of centriole elimination

We performed CLEM to characterize the centriole elimination process at the ultrastructural level to gain insights into the mechanisms leading to organelle removal. We dissected, fixed and imaged two gonads expressing GFP::SAS-7, followed by resin embedding and 50 nm serial section EM analysis. A total of 69 foci bearing GFP::SAS-7 were analyzed in this manner, representing successive stages of oogenesis and, therefore, centriole elimination. Each nucleus with an accompanying GFP::SAS-7 focus was first visualized by brightfield and then identified in the corresponding serial sections, thus providing spatial coordinates to spot centrioles (Fig. 5A-5C). As previously reported (Mikeladze-Dvali et al., 2012; Woglar et al., 2022), we found that nuclei in early meiotic prophase possess four centriolar units, two mature centrioles and two associated procentrioles (Fig. 5D). Distinct ultrastructural elements can be observed in top views at this stage, including the inner-most tube, the central tube, microtubule singlets, and the peripheral-most paddlewheel (Fig. 5E) (Sugioka et al., 2017; Woglar et al., 2022). Strikingly, during diplotene, which corresponds approximately to when the intensity of the GFP::SAS-7 focus begins to decline (see Fig. 1), ultrastructural analysis revealed that the central tube is no longer detectable (Fig. 5F). In addition, this is accompanied by an increase of organelle diameter (Fig. 5I, 5J). Loss of centriole integrity was observed thereafter, with microtubule singlets no longer being recognized (Fig. 5G), which was followed by the complete loss of a detectable centriole (Fig. 5H). Interestingly, this is the case despite GFP::SAS-7 remaining present in a focus at this stage, which will be cleared eventually as reported above.

**Figure 5.**
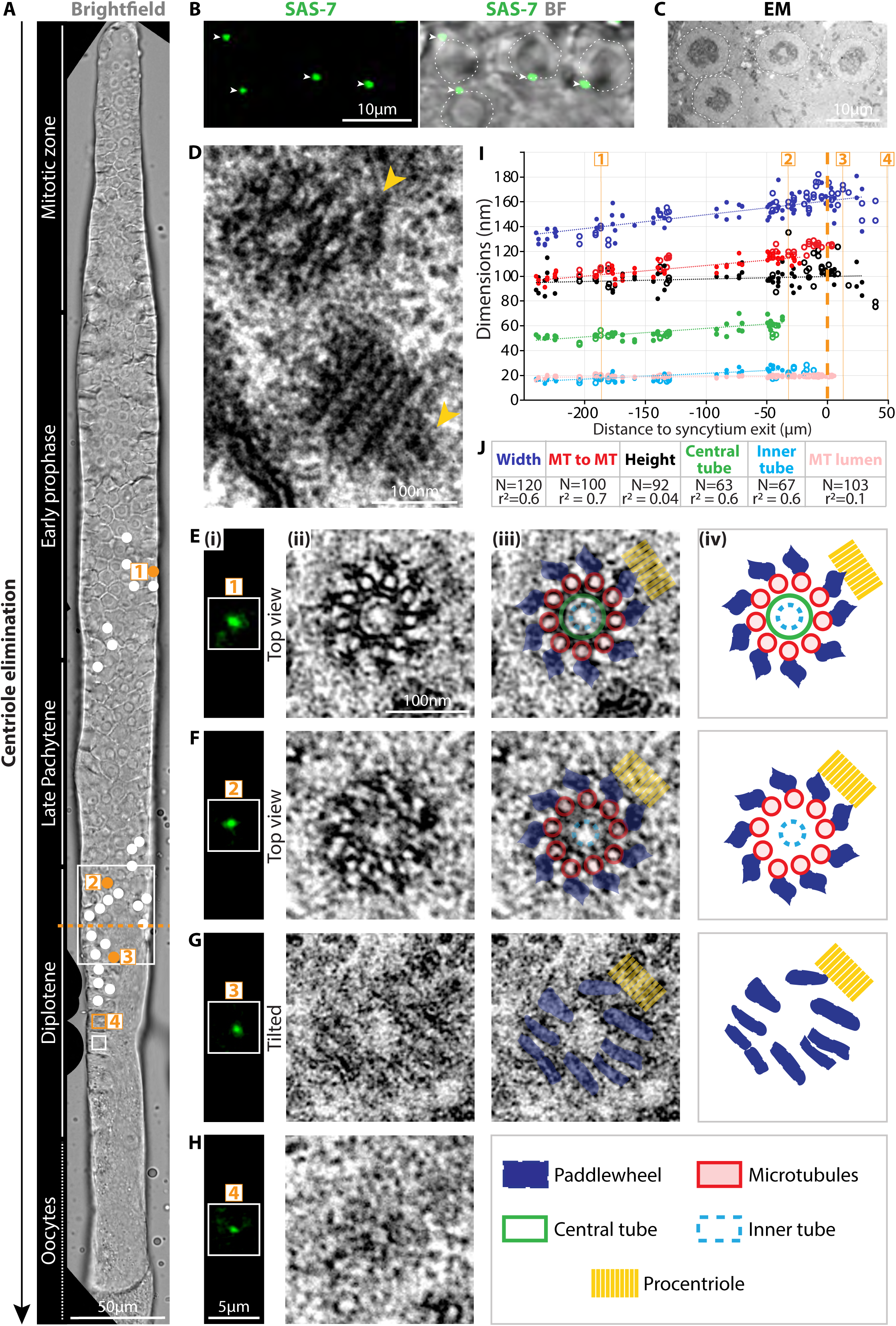
Ultrastructural analysis reveals loss of central tube and centriole widening at elimination onset. **A.** Single focal plane brightfield image from a gonad expressing GFP::SAS-7 imaged after chemical fixation and before resin embedding for electron-microscopy (EM) serial sectioning. The gonad was straightened computationally for analysis purposes. Serially-sectioned nuclei with centrioles are marked with filled white disks, those without centrioles with empty squares. Orange numbers correspond to exemplary cases shown in E-H. Dashed orange line marks the border between the syncytial region (to the top) and the cellularized region (to the bottom). White rectangle corresponds to inset in B. **B.** Magnified inset from A showing centriolar GFP::SAS-7 (left, arrowheads) relative to nuclei (right, Brightfield (BF), dashed circles). **C.** Corresponding 50nm EM section. Note alignment of nuclei between B and C (dashed circles), which helps locating region corresponding to GFP::SAS-7 foci in the EM sections. **D.** 50nm EM section of a pair of centrioles, one viewed from the top (top) and one from the side (bottom), each with a procentriole (orange arrowheads). **E-H. (i)** GFP::SAS-7 foci from the four positions numbered in orange in A. **(ii)** Corresponding 50nm EM sections of centrioles in top or tilted views, as indicated, illustrating stages of elimination, with loss of central tube (F), of microtubule singlets (G), and of recognizable centriolar ultrastructure (H). **(iii)** Overlay of EM images with indicated ultrastructural compartments. **(iv)** Schematic representation of successive stages of elimination. The electron dense material at the bottom right of images E(ii) and E(iii) is part of the endoplasmic reticulum. **I.** Centriole width and height, as well as diameter of indicated ultrastructural compartments, at successive stages of centriole elimination, using distance from syncytium exit (dashed orange line) as a proxy for time. Data acquired from 2 independent gonads (filled and empty circles, respectively), using top, longitudinal and slightly tilted views for analysis (Materials and methods). Numbered orange vertical lines indicate measurements made on centrioles shown in E, F, G and H. **J**. Color code and number of data points for each series in I, as well as r^2^ values for trendlines depicted as dashed lines. MT: microtubules.

Taken together, these results reveal that centriole elimination during *C. elegans* oogenesis is characterized by an initial loss of organelle integrity, followed by loss of an amorphous cluster of centrosomal proteins. Moreover, these findings raise the possibility that the first observable ultrastructural alteration, namely central tube removal, may cause subsequent loss of centriole integrity.

### U-Ex-STED reveals that SAS-1 leaves centrioles at the onset of centriole elimination

U-Ex-STED analysis of gonad centrioles established that the central tube harbors the microtubule-binding protein SAS-1 (Woglar et al., 2022). Suggestively, SAS-1 is required for the stability of sperm-derived centrioles in the zygote and of centrioles assembled subsequently during embryogenesis (von Tobel et al., 2014). Together with our finding that central tube loss is a hallmark of centriole elimination onset during oogenesis, this raises the possibility that SAS-1 removal may mark the first step of organelle removal in the female germline. To investigate this possibility, we used U-Ex-STED to analyze the distribution of endogenously tagged SAS-1::3xFLAG during the course of the changes uncovered by EM, in relationship to SAS-7 and SAS-4 distribution. Since the central tube is the first ultrastructural element to vanish, we expected fluorescence intensity of the SAS-1 focus to diminish before that of SAS-7 and SAS-4, and found this to be the case indeed (Fig. 6A-6C). We used U-Ex-STED to refine the timing of SAS-1 loss relative to that of microtubules, finding that GBP::RFP::SAS-1 fluorescence at centrioles decays faster than that of microtubules, consistent with the early loss of the central tube observed by EM (Fig. 6D, 6G). Using an analogous experimental strategy, we set out to investigate the integrity of other ultrastructural elements in relationship to microtubules. Importantly, whereas SPD-5 rings are dismantled concomitantly with microtubule singlets (Fig. S4A, S4B), consistent with the early loss of SPD-5 uncovered in live specimens (see Fig. 1I, 1J), we found that centriolar microtubules are lost before SPD-2 or SAS-4 (Fig. 6E-6G). This finding likely explains why foci containing centriolar proteins with decaying signal intensity remain present in maturing oocytes despite the complete loss of ultrastructural integrity observed by EM (Fig. 5H). Moreover, like in the EM analysis, U-Ex-STED uncovered a widening of centrioles as they progress through meiosis, as evidenced by measurements of SPD-2, microtubules and SAS-4 ring diameters (Fig. 5H).

**Figure 6.**
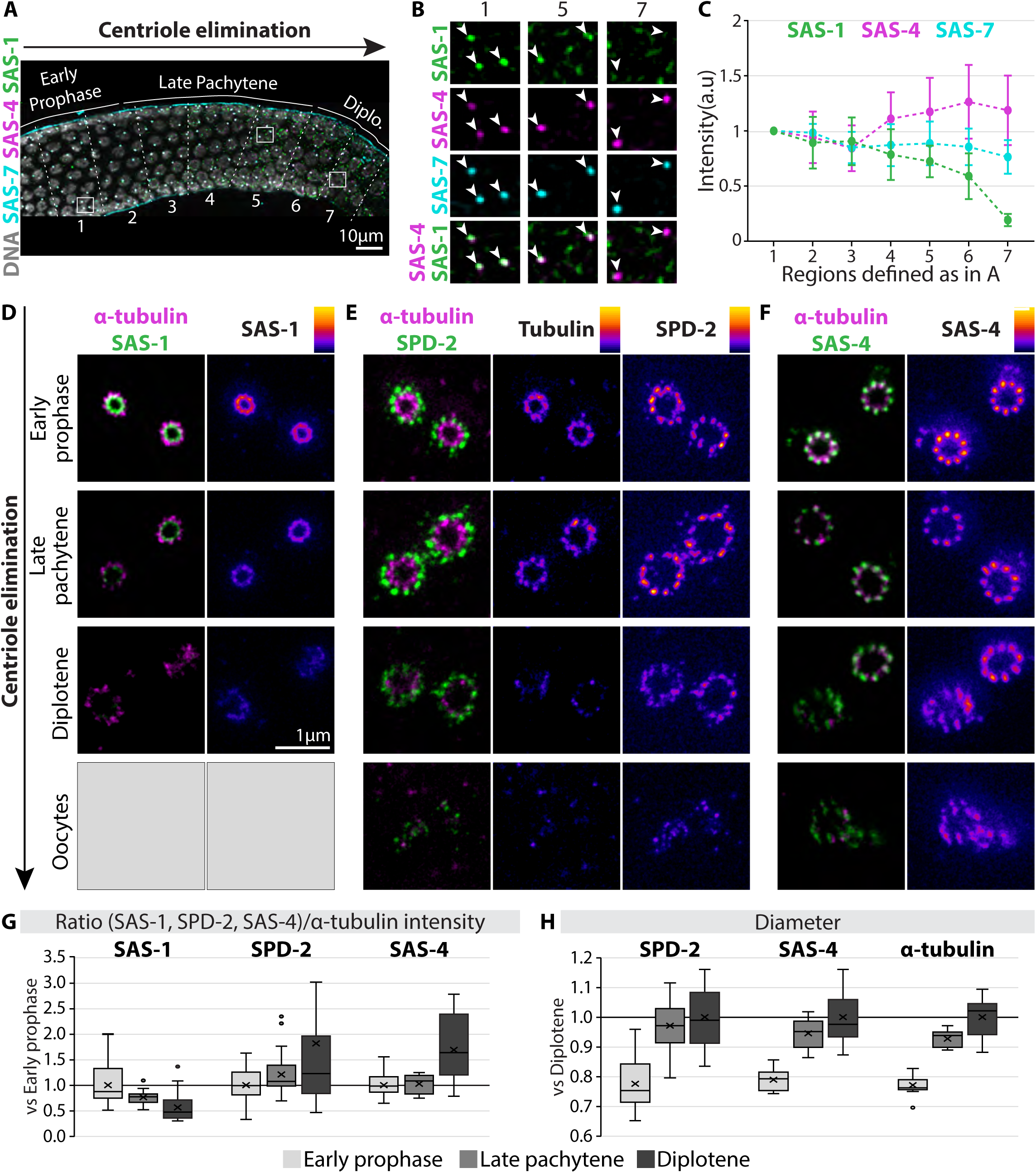
Centriole widening occurs concomitantly with the loss of the central tube protein SAS-1 at the onset of centriole elimination. **A.** Gonad immunostained with antibodies recognizing SAS-1::3xFLAG (green), SAS-4 (magenta) and GFP::SAS-7 (cyan); DNA counterstain: grey. Seven sub-regions within early prophase, late pachytene and diplotene regions were defined as indicated. Each region is approximately three times the diameter of a diplotene nucleus. White squares correspond to insets in B. **B.** Magnified insets from A, illustrating earlier loss of centriolar SAS-1 compared to SAS-7 and SAS-4 (arrowheads). **C.** Normalized fluorescence intensity for SAS-1, SAS-7 and SAS-4 across the seven sub-regions illustrated in panel A. The background-subtracted signal of foci was averaged for each region and normalized to the mean value in region 1. Mean ± SD for each region; N= 7 gonads. **D-F.** U-Ex-STED of centrioles at indicated stages (early prophase, late pachytene and diplotene) from worms expressing GBP::RFP::SAS-1 (D), SPD-2::GFP (E) and SAS-4 (F), stained for RFP or GFP and co-stained with an antibody against α-tubulin. The LUT “Fire” reports fluorescence intensity (from bright to dim: white, yellow, red, purple, blue and black). The expansion factor is the same for each sample so that relative sizes within each series can be compared (i.e. within D, E and F). Note that we used GBP::RFP::SAS-1 instead of SAS-1::3xFLAG in these experiments as RFP antibodies gave stronger signals than FLAG antibodies, allowing better detection following U-Ex-STED. D, grey boxes were used because of the impossibility to identify centrioles remnant based on the tubulin staining at the oocytes stage. **G.** Normalized fluorescence intensity of GBP::RFP::SAS-1, SPD-2::GFP and SAS-4 during prophase progression determined from U-Ex-STED images. All values were background-subtracted and first normalized to the α-tubulin signal intensity in the same centriole and then to the mean value of early prophase within each series. N (early prophase, late pachytene, diplotene): GBP::RFP::SAS-1/α-tubulin: 15, 14, 16; SPD-2::GFP/α-tubulin: 42, 26, 41; SAS-4/α-tubulin: 22, 10, 15. In box-plots in this and other figures, the middle line corresponds to the median and the cross to the mean; boxes include 50% of values (IQR) and whiskers show the range of values within 1.5*IQR. **H.** Relative diameter during prophase progression of rings formed by the three indicated proteins from centriole top views imaged by U-Ex-STED. For each series, values were normalized to the mean diameter at diplotene. N (early prophase, late pachytene, diplotene): SPD-2: 34, 26, 31; SAS-4: 22, 10, 18; α-tubulin: 17, 10, 16.

Together, these data demonstrate that removal of the microtubule-binding protein SAS-1 is an early event in the sequence of events leading to organelle demise.

### SAS-1 is required for centriole structural integrity and stability during oogenesis

If SAS-1 departure from centrioles not only marks the onset of oogenesis centriole elimination, but is also important for this process to occur, then impairing *sas-1* function should exacerbate loss of centriole integrity. To test this prediction, we performed U-Ex-STED on gonads from the strong reduction-of-function allele *sas-1(t1521ts)* at the restrictive temperature. Strikingly, we found that centrioles are as wide in early prophase as they are in diplotene in this mutant background, whereas widening only occurs during diplotene in the control (Fig. 7A-7C). Moreover, whereas SAS-4 is normally present in approximately equal amounts next to each of the nine centriolar microtubules in early prophase, this is not the case in *sas-1(t1521)* mutant animals, where intensities are more variable (Fig. 7D). Together, these two data sets suggest that centriole integrity is already affected in early prophase when *sas-1* function is compromised. Furthermore, U-Ex-STED analysis revealed that centriolar microtubules and SAS-4 signals decay faster during meiosis progression in *sas-1(t1521ts)* mutant animals than in the control (Fig. 7E, 7F). This is accompanied by premature loss of organelle integrity, as evidenced by the fraction of centrioles in diplotene with less than nine foci of centriolar SAS-4 and completely disorganized centrioles (Fig. 6G). Overall, these data demonstrate that SAS-1 is required for centriole stability during oogenesis, and suggest that SAS-1 departure from centrioles may trigger organelle elimination from the female germ line.

**Figure 7.**
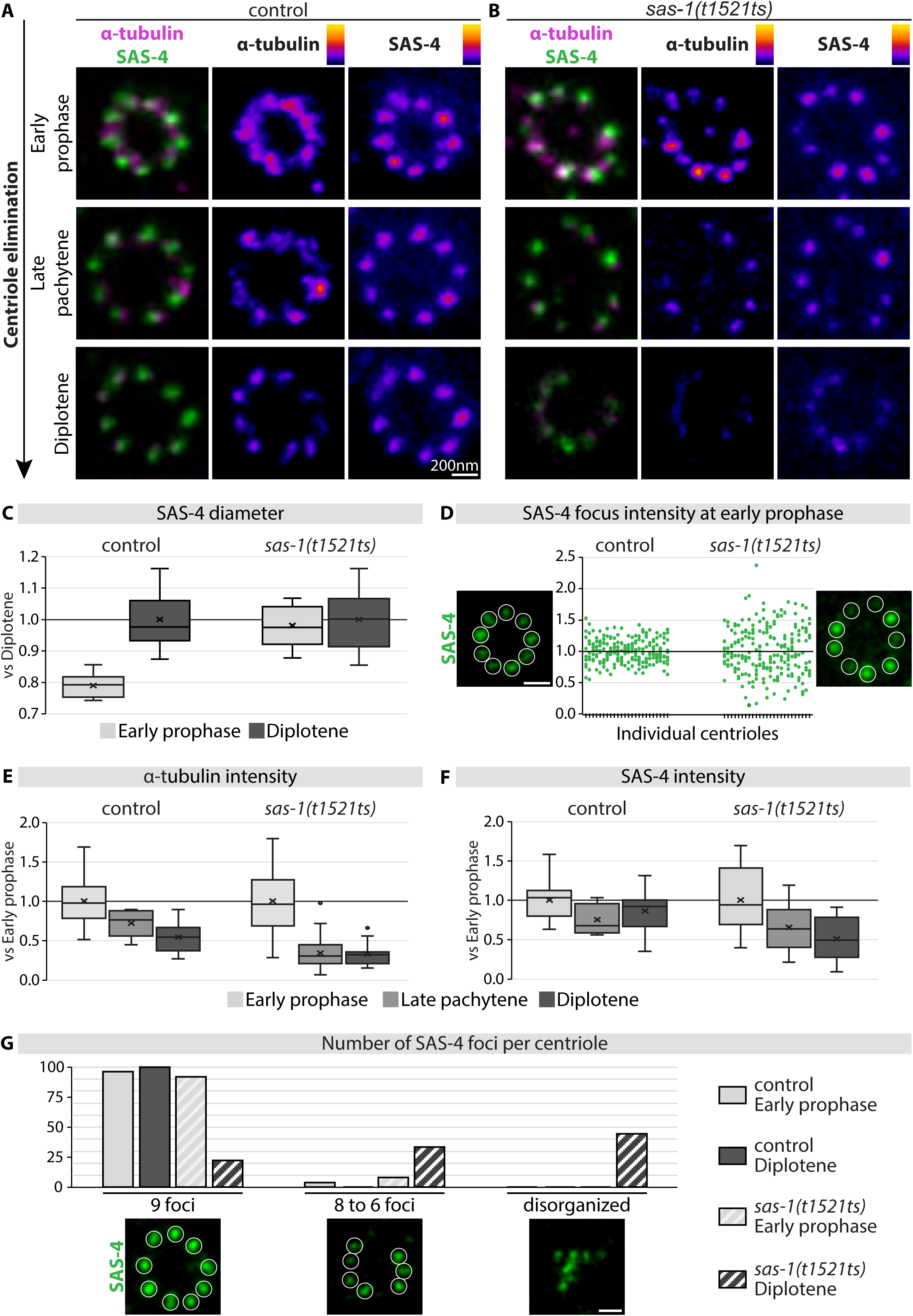
SAS-1 is required for proper centriole integrity during oogenesis. **A, B.** U-Ex-STED top views of centrioles during prophase progression from control (A) and *sas-1(t1521ts)* mutant at the restrictive temperature (B) stained for α-tubulin (magenta) and SAS-4 (green). **C.** Corresponding SAS-4 relative ring diameter from top views imaged by U-Ex-STED. For each series, values were normalized by the mean diameter at diplotene. N (early prophase, diplotene): control: 21, 18; *sas-1(t1521ts)*: 22, 13. **D.** U-Ex-STED top views of early prophase centrioles from control (left) and *sas-1(t1521ts)* mutant (right), together with corresponding brightness distributions of the 9 discrete SAS-4 signals (center). Each vertical column displays the 9 signals measured in one centriole normalized to the mean signal intensity of that centriole. The two distributions are statistically different (Levene test, p-value<0.00001). **E, F.** Evolution of normalized α-tubulin (E) and SAS-4 (F) fluorescence intensity during prophase progression in samples imaged by U-Ex-STED. Values were background-subtracted and normalized to the mean of early prophase measurements in each series. N (prophase, late pachytene, diplotene): control: 17, 10, 16; *sas-1(t1521ts)*: 24, 32, 20. **G.** Number of discrete SAS-4 foci in centrioles in early prophase and diplotene nuclei of control and *sas-1(t1521ts)* mutant, together with exemplary U-Ex-STED images. N (early prophase, diplotene): control: 27, 14; *sas-1(t1521ts)*: 25, 18.

## DISCUSSION

We established with precision the course of events leading to centriole elimination during *C. elegans* oogenesis using multi-scale microscopy and molecular genetic approaches. We discovered that oogenesis centriole elimination is characterized by an initial loss of the central tube and organelle integrity, followed by the gradual disappearance of an amorphous cluster of centrosomal proteins. The latter often occurs after detachment from the nucleus and movement to the plasma membrane in a microtubule- and dynein-dependent manner, although such movement is not essential for disappearance. Finally, we demonstrate that the central tube protein SAS-1 is the first component to leave centrioles and propose that this event triggers centriole elimination.

### Clearance of amorphous cluster of centrosomal proteins

We found that, following the loss of centrioles recognizable by EM, an amorphous cluster containing several centrosomal proteins persists up to the −2 oocyte, later than what was previously reported (Matsuura et al., 2016; Mikeladze-Dvali et al., 2012). The prolonged detection of foci in the present study probably reflects the use of brighter CRISPR/Cas9-based and transgenic reporters, as well as more sensitive microscopy. Using live imaging, we show that in ∼80% of cases, cluster removal occurs after detachment from the nucleus and movement to the plasma membrane in a microtubule- and dynein-dependent manner. Whereas the centrosome is known to be linked to the nucleus in the embryo via ZYG-12/SUN-1 (Malone et al., 2003; Zhou et al., 2009), the components mediating the analogous link in the gonad are not known. Our findings suggest that SPD-5 and the PCM are involved. Indeed, in the absence of these components upon *pcmd-1* inactivation, centrioles tend to detach from nuclei already in early prophase, localizing to the rachis and the loop region. Moreover, in the control condition, the SPD-5 focus is reduced to ∼30% of initial levels already in −7 oocytes, shortly before cluster detachment initiates, compatible with a causative link. Alternatively, a more global remodeling of the nuclear envelope might promote detachment of different molecular complexes in maturing oocytes since P-granules, which do not contain PCM proteins, also detach from the nuclear envelope around that time (Spike et al., 2008).

Cluster movement to the plasma membrane occurs at an average velocity of ∼0.65μm/min, which is very slow considering that it relies on microtubules and dynein, and partly on kinesin. Dynein and kinesin motor proteins in *C. elegans* exhibit velocities along microtubules that are two orders of magnitude faster (Gönczy et al., 1999; Pierce et al., 1999). Interestingly, centrioles also translocate in a microtubule- and dynein-dependent manner from the nuclear vicinity to the tip of the dendrite in the PQR sensory neuron in *C. elegans*, a movement that is accompanied by gradual loss of centriolar proteins (Li et al., 2017). Such movement was proposed to result from transient interactions between dynein and centrioles (Li et al., 2017), and it may be that the same mechanism is at play during oogenesis. Alternatively, given that the microtubule network is nucleated mainly from the nuclear envelope and the plasma membrane in *C. elegans* oocytes, with dynein being required for overall network organization (Zhou et al., 2009), not having these components may lead to a disorganized cellular state incompatible with cluster movement. Regardless, importantly, we show that cluster movement to the plasma membrane is not essential for disappearance of clustered centrosomal proteins, since ∼20% of clusters remain in the vicinity of the nucleus and yet are disposed of. Why should such a clearance step be needed when centriole integrity has been lost already? It has been established notably in human cells that centriolar protein assemblies that are not *bona fide* organelles, in that they do not harbor centriolar microtubules, can nevertheless act as MTOCs, and thereby lead to faulty spindle assembly (Balestra et al., 2021; Li et al., 2012; Shiratsuchi et al., 2015). Therefore, clearance of the cluster of centrosomal proteins in *C. elegans* may serve as a safety mechanism to prevent regaining MTOC activity in the oocyte and the zygote.

### Diversity in Polo-like kinase requirement for oogenesis centriole elimination

Whereas Polo kinase and the PCM play a critical role in centriole elimination during *Drosophila* oogenesis (Pimenta-Marques et al., 2016), we demonstrate that the situation differs in *C. elegans*. Indeed, single or combined depletion of PLKs, as well as removal of the PCM core component SPD-5, do not have a discernable impact on the timing of centriole elimination in the worm. Whereas time-resolved analysis by serial-section EM would be needed to ascertain that no change in centriole ultrastructure has occurred, several considerations may explain the difference between the two systems. First, whereas Polo is present at centrioles in the *Drosophila* germ line in ∼40% of stages 7/8 oocytes and in ∼15% of stage 9/12 oocytes, when elimination occurs, PLK-1 and PLK-2 are located at centrioles only in proliferating germ cells in *C. elegans*, much before elimination begins (Jaramillo-Lambert et al., 2007; Mikeladze-Dvali et al., 2012). Second, whereas Polo is required to maintain the PCM in fly oocytes, we show here that depleting PLKs in the worm has no impact on the timing of loss of the PCM core protein SPD-5. Third, we note that 64 centrioles, 60 from the 15 nurse cells plus 4 from the oocyte proper, end up clustered in the *Drosophila* oocyte(Pimenta-Marques et al., 2016), whereas merely 4 centrioles are present in the vicinity of each germ nucleus in the *C. elegans* gonad. Perhaps a large PCM surrounding a cluster of 64 centrioles prevents the elimination machinery to reach centrioles in *Drosophila*, such that prior PCM disassembly is a prerequisite for organelle removal. Inhibiting Plk1 in starfish oocytes does not alter disappearance timing of foci bearing the centriolar marker Centrin2, suggesting that also in this system centriole elimination is not regulated by this kinase (Pierron et al., 2020). Future work should help clarify to what extent the requirement in *Drosophila* oogenesis is deployed in other centriole elimination settings.

### Central tube and SAS-1 removal mark the onset of centriole elimination

By combining EM and U-Ex-STED, we reveal that the first detectable sign of centriole elimination is the disappearance of the central tube, together with a widening of the centriole, which precede the loss of microtubule singlets and of organelle integrity. These observations are partially reminiscent of the centriole degeneration steps at the time of cilium formation in amphid neurons, with the loss of the inner core structure and the splaying of microtubules (Doroquez et al., 2014; Nechipurenko et al., 2017; Serwas et al., 2017). Moreover, the ∼40nm increase of organelle diameter measured during that degeneration process is in the range of that determined here during oogenesis centriole elimination (Nechipurenko et al., 2017). However, as opposed to the early loss of SAS-6 and then SAS-4 in amphid neurons (Nechipurenko et al., 2017; Serwas et al., 2017), we find here that these proteins persist until the end of centriole elimination during oogenesis.

Our analysis uncovered a critical role for SAS-1 during oogenesis centriole elimination. We find that SAS-1 is the earliest component analyzed to depart from centrioles during this process, in line it being a central tube component (Woglar et al., 2022). In addition, U-Ex-STED analysis revealed that centrioles are as wide during early prophase as they are in diplotene in *sas-1(t1521ts)* mutant animals. Moreover, centrioles in *sas-1(t1521ts)* mutants are eliminated faster than in control conditions. Together, these findings define SAS-1 as a key player in the initiation of centriole elimination.

SAS-1 was known already to be required for the stability of sperm-derived centrioles: centrioles derived from *sas-1* mutant sperm lose their integrity shortly after fertilization, as evidenced by the reduction of centriolar fluorescent markers intensity and ultrastructural analysis (von Tobel et al., 2014). Moreover, centrioles assembled in embryos with compromised *sas-1* function are likewise unstable (von Tobel et al., 2014). Interestingly, SAS-1 associates with microtubules when expressed in human cells (von Tobel et al., 2014). Combined with the U-Ex-STED data, these findings taken together suggest that the central tube, and SAS-1 in particular, may act as an inner brace holding the microtubule singlets together.

SAS-1 is an ortholog of the human ciliopathy protein C2CD3 (C2 domain containing 3), which is involved in human centriole length control and organization of distal appendages, where it forms a distal brace perhaps analogous to the worm central tube (Chang et al., 2023; Gaudin et al., 2022; Thauvin-robinet et al., 2014). *C. elegans* centrioles lack apparent proximal/distal polarity, with all constituent proteins known to date being homogeneously distributed along its length. Whether C2CD3 plays a role in centriole stability in human cells or may be targeted for disassembly in specific cellular context remains to be tested. Because centrioles disassemble earlier in *sas-1* mutant without impacting oogenesis, a potential involvement of C2CD3 in vertebrate oogenesis centriole elimination may have gone undetected.

To conclude, our findings contribute to the understanding of centriole elimination during oogenesis, a fundamental process that is essential for the sexual reproduction of metazoan organisms. Furthermore, we anticipate the sequence of events unraveled here, with the key role of SAS-1 at the onset of the elimination process, to serve as a platform to understand centriole fate modulation in other physiological and pathological contexts.

## ACKNOWLEDGEMENTS

We are grateful to the laboratories of Bruce Bowerman, Alex Dammermann, Jessica Feldman, Anthony Hyman, Kevin O’Connell and Anne M. Villeneuve for their gift of worm strains. Some other strains were provided by the *Caenorhabditis* Genetics Center (CGC), which is funded by the NIH Office of Research Infrastructure Programs (P40 OD010440). We thank Graham Knott, head of the BIO-EM platform, as well as Simon Berger for training with the microfluidic chamber and providing the tools. We are grateful to Gabriela Garcia Rodriguez and Nils Kalbfuss for comments on the manuscript, as well as to Léo Burgy for help with statistical analysis. This work was funded by post-doctoral fellowships from EMBO to M.P. (ALTF 1426-2016), the European Union to A. W. (MCSA-IF 588594) and DFG to T. M.-D. (MI1867/1-3), as well as by grants to P.G. from the Novartis Foundation for Medical-Biological Research (18C186) and the Swiss National Science Foundation (310030_197749).

## AUTHOR CONTRIBUTIONS

Conceptualization, M.P., A.W., T.M-D., P.G.; Investigation, M.P., A.W., C.B., K.J., T. M-D.; Writing – Original Draft, M.P., A.W., K.J. and P.G.; Funding Acquisition, M.P., A.W., P.G., T.M-D; Supervision, P.G.

## Materials and methods

### *C. elegans* strains and RNAi

The *C. elegans* lines generated for this study (see Table1) are available from the lead contact upon request. Strains were maintained following standard methods on nematode growth medium (NGM) plates seeded with *Escherichia coli* OP50 as food source (Brenner, 1974). Strains were kept at 20°C or 24°C, except for thermosensitive strains, which were kept at 16°C until the L4 stage, when they were shifted to 24°C or 25°C for 20-24 h prior to imaging. The list of strains used in this study is given in Table S1. Synchronized populations were obtained by allowing 20-50 gravid adults to lay eggs for 1 h at room temperature (RT) and imaging gonads of the resulting adults after ∼65h of growth at 20°C. RNAi by feeding was performed with clones from either Ahringer or Vidal library to deplete *plk-1*(Vidal), *tba-2* (Vidal), *dhc-1* (Arhinger), *dlc-1* (Vidal), *act-1* (Vidal) and *unc-116* (Arhinger), feeding L3/young L4s at 24°C and imaging 20-24h thereafter.

### Gonad spreading

Spreading of *C. elegans* gonads was performed similarly as in (Woglar et al., 2022). Gonads of ∼ 1000 adult worms were dissected in 30 μL PBS-T (0.2 x PBS, 1:1000 Tween 20) on an ethanol-washed 22×40 mm coverslip. 5-10 μL of dissected gonads were then pipetted onto a new ethanol-washed 22×40 mm coverslip and 50 μL of spreading solution (for one coverslip, 50 μL: 32 μL of Fixative was added (4% w/v Paraformaldehyde and 3.2% w/v Sucrose in water), 16 μL of Lipsol solution (1% v/v/ Lipsol in water), 2 μL of Sarcosyl solution (1% w/v of Sarcosyl in water), and gonads were immediately distributed over the coverslip using a pipette tip. Coverslips were left to dry at room temperature (RT) followed by incubation at 37°C for 1h. Coverslips were either processed for staining and expansion or stored at −80°C.

### Gonads expansion for U-Ex-STED

Ultrastructure expansion microscopy was performed as in (Woglar et al., 2022). In brief, dried coverslips were incubated for 20 min in methanol at −20°C and washed 3 times in PBS-T for 5 min, followed by two 5 min washes in PBS. Coverslips were incubated with mild agitation in a 5cm Petri dish overnight at RT in Acrylamide/Formaldehyde solution (1% Acrylamide and 1% Formaldehyde in PBS). Thereafter, coverslips were washed 3 times 5 min in PBS. For gelation, coverslips were incubated in 50 µl monomer solution (19% (wt/wt) Sodium Acrylate, 10% (wt/wt) Acrylamide, 0.05% (wt/wt) BIS in PBS) supplemented with 0.5% Tetramethylethylenediamine (TEMED) and 0.5% Amonium Persulfate (APS) on a piece of Parafilm for 1h at 37°C in a moist chamber in the dark. All subsequent steps were carried out under mild agitation at room temperature unless otherwise stated. Gels were incubated in 5cm Petri dishes for 15 min in denaturation buffer (200 mM SDS, 200 mM NaCl and 50 mM Tris in distilled water, pH=9) followed by incubation for 1 h on a 95°C hot plate in fresh denaturation buffer. Gels were transferred to 15 cm Petri dishes, washed with distilled water 5 times for 20 min, followed by incubation in distilled water overnight at 4°C. The expansion factor was estimated by measuring the gel size with a ruler.

### Immunostainings on whole dissected gonads and expanded germ cell nuclei

Gonads were dissected in sperm buffer (50 mM Hepes (pH 7.0), 50 mM NACL, 25 mM KCL, 5 mM CaCl2, 1 mM MgSO4, 50 mM Glucose, 1 mg/ml BSA), transferred onto slides (Marienfeld, 1000200) coated with poly-lysine (2 mg/ml in PBS), freeze-cracked and fixed in cold methanol for 5 min. Slides were then incubated overnight at RT in blocking buffer (3% BSA in PBST) with primary antibodies as follows: mouse anti-IFA-1 (1:100, ATCC-TIB-131 (Pruss et al., 1981)); chicken-anti-GFP (1:500, Abcam ab13970), rabbit-anti-SAS-4 (1:800 (Leidel and Gönczy, 2003)); mouse-anti-Flag (1:200, Sigma-Aldrich F1804). Slides were then washed twice 10 min in PBS-T, incubated with anti-chicken Alexa Fluor 488 (1:1000, A11039), anti-rabbit Alexa Fluor 647 (1:1000, A10523), anti-mouse Alexa Fluor 568 (1:1000, A11004) in PBS-T for 2h at RT, washed three times 10min in PBS-T, counterstained with Hoechst (1:1000) and mounted in mounting medium (4% n-Propyl-Gallate, 90% Glycerol, 1xPBS).

After expansion, gels were cut into pieces fitting into a 5 cm Petri dish. Prior to staining, gels were blocked for 1h at RT in blocking buffer (10mM HEPES (pH=7.4), 3% BSA, 0.1% Tween 20, sodium azide (0.05%)), followed by incubation overnight at RT with primary antibodies diluted in blocking buffer. Gels were washed three times in blocking buffer for 10 min each, before incubation with secondary antibodies diluted in blocking buffer supplemented with 0.7 ug/L Hoechst for 3 h at 37°C in the dark. Gels were washed three times in blocking buffer for 10 min each before transfer into a 10cm Petri dish for re-expansion by washing 6 times 20 min in distilled water. For imaging, gels were cut and mounted on a 60×24 mm coverslip coated with poly-D-lysine (Sigma, # P1024) diluted in water (2 mg/ml) and supported on both longitudinal sides with capillaries attached with superglue. To prevent drying, the edges of the gel were covered with VaLaP (1:1:1 petroleum:jelly:lanolin:paraffin wax), and the gel covered with Halocarbon oil 700 for imaging. Antibodies used on expanded gels were as follows: rabbit anti-SAS-6 (1:1000 (Leidel et al., 2005)), rabbit anti-SAS-4 (1:800 (Leidel and Gönczy, 2003)), rabbit anti-SAS-5 (1 :50 (Delattre et al., 2004)), rabbit anti-GFP (1:250, a gift from Viesturs Simanis), mouse anti-RFP(RF5R) (1:500, Thermo Fisher, MA5-15257), rabbit anti-α-tubulin (1:500, Abcam (ab52866)) and rat anti-tyrosine α-tubulin (1:500, EMD Millipore, MAB1864). Secondary antibodies (all used at 1:1000) were as follows: donkey anti-rabbit conjugated to Alexa Fluor 594 (Abcam, ab150072), goat anti-rat conjugated to Alexa Fluor 594 (Thermo Fisher, A11007), goat anti-rabbit conjugated to Alexa Fluor 488 (Thermo Fisher, A11034) and donkey anti-rat conjugated to Alexa Fluor 488 (Invitrogen, A21208).

### Microscopy

For live gonad imaging, worms were mounted on 2% agarose pads and immobilized with 8 μl of 100mM sodium azide (NaN_3_). Spinning disc confocal imaging was performed using an inverted Olympus IX 83 motorized microscope equipped with a Yokogawa spinning disk CSU-W1 head, a 60× (NA 1.42 U PLAN S APO) objective, an ImagEMX2 EMCCD and an Orca Flash 4.0 sCMOS camera, controlled by VisiView software. Widefield imaging was performed using an inverted Nikon eclipse Ti2-U widefield microscope, with a Nikon Plan Apo 60x/1.40 ∞/0.17 WD 0.13 objective. The microscope was equipped with a Nano Z500 super long-range Piezo z-stage, a Märzhäuser Tango 3 desktop controller together with the Ergodrive 3 3-axes operating device, a Lumencor Spectra4 as light source, an Andor Zyla-4.2P CL10 sCMOS camera, and was controlled by Micromanager (Edelstein et al., 2014). For time-lapse imaging, worms were partially immobilized in a microfluidic device enabling continuous feeding (Berger et al., 2018). For U-Ex-STED, stages of meiotic prophase were identified by chromatin morphology visualized with Hoechst (Phillips et al., 2009). 2D-STED images were acquired on a Leica TCS SP8 STED 3X microscope with a 100 x 1.4 NA oil-immersion objective, using 488 nm and 589 nm excitation, and 592 nm and 775 nm pulsed lasers for depletion. 1-pixel Gaussian blur was applied to all images for analysis and display. For display, brightness and contrast was adjusted in the individual channels using Fiji, keeping the same settings within a series.

### Image processing and analysis

Microscopy images were stitched, rotated, z-projected; gray levels were then adjusted using Fiji (ImageJ). Gonad images and insets are maximum intensity projection of relevant planes. For dual color quantifications of live imaging at the spinning disc confocal, a ROI of 15×15 pixels was centered on each focus at the z-slice displaying the maximum brightness; Max intensity projection of 12 μm around this position was applied to generate a 2D image. Projection on one of the axes then provided a 1D intensity profile. We fitted the function y=a+b*x+c*exp(-/x-d)*(x-d)/e) on this profile and subtracted linear background through the first and the last data point. The integration over the resulting curve gave the reported intensity value for one focus. Values for foci in oocytes from different gonads located at the same position with respect to the spermatheca were averaged. Fluorescence intensity curves were normalized according to the maximal and minimal values of all oocytes for each marker. ImageJ macro used for quantifications is on GitHub (https://github.com/UPGON/pierron2023-centriole-elimination). In the case of RME-2::GFP, an oocyte was scored as positive when the four sides of the plasma membrane exhibited GFP fluorescence. For quantification of immunostaining in Fig. 6C, sum-projected images of gonads were divided into seven regions separated approximately by three times the diameter of diplotene nuclei at the beginning of cellularization. ROIs of 0.5-2 μm_2_ were drawn around each focus, as well as just next to it for background subtraction. The background-corrected signal was then determined for each channel, and the average signal for each region normalized with that of the most-distal region 1.

### Timelapse imaging and movie analysis

For long-term live imaging, worms were loaded and immobilized in the microfluidic as described (Berger et al., 2018). Briefly, synchronized day 1 adults were collected and washed 3 times in fresh S-basal buffer (5.85 g NaCl, 1 g K_2_HPO_4_, 6 g KH_2_PO, 1 ml cholesterol (5 mg/ml in ethanol), H_2_O to 1L) and left to sediment. In parallel, a 50 ml Falcon tube of NA22 grown overnight at 37°C was centrifuged (at 4000rcf for 20 min) to obtain a bacteria pellet. This pellet was washed 3 times in S-basal and resuspended in 1 ml S-basal supplemented with 0.65mL of Optiprep (density matching to prevent bacteria from segregating, 1114542 by Serumwerk Bernburg for Alere Technologies) and 0.332 mL of S-Basal+1% Pluronic F127 (P2443, Sigma-Aldrich). This food mixture was filtered through a 10μm cell strainer (#43-10010-50, pluriSelect) to remove bacteria clumps. The microfluidic device was set up on the Nikon eclipse Ti2-U widefield microscope. The food supply was connected to the device, and the worms were loaded and immobilized for imaging. Worms were positioned so that the proximal part of the posterior gonad was always closest to the coverslip. Movies were acquired during of 3-5 h with 2min intervals and binning 2. We filmed 13 worms among which 2 were excluded because of apparent signs of starvation. Among the 11 worms analyzed, 6 are expressing MEL-28::GFP and 5 are expressing RME-2::GFP, together with RFP::SAS-7. We focused on oocytes in positions −6 to −4 at the onset of the experiment, so that the entire movement of foci could be monitored. Cropped images were extracted and aligned based on the center of the nucleus, which was manually selected at each timepoint for a given oocyte. using. For 7 movies, kymographs were generated using a line width of 40 and the KymographBuilder plugin in Fiji (Mary et al., 2016). Centriolar foci remnant speed was calculated using the maximum distance between the green signal and the magenta signal on the Kymograph during the centriolar focus migration phase. The graph was generated using PlotsofData (Postma and Goedhart, 2019).

### CLEM analysis

Gonads of genotype sas-7(or1940[gfp::sas-7])III; glo-1(zu931)X; itIs37[pie-1p::mCherry::H2B, unc-119(+)] or ltSi202[pVV103/ pOD1021; Pspd-2::GFP::SPD-5 RNAiresistant;cb-unc-119(+)]II; sas-7(is1[tagRFP::sas-7+loxP])III; glo-1(zu931)X were dissected in sperm buffer and transferred onto poly-lysine-coated MatTek glass bottom dishes. 3D imaging of gonads was performed using the previously mentioned Nikon widefield microscope set up, before and after an approximately 2h30 min fixation at RT in 2.2% glutaraldehyde, 0.9% Paraformaldehyde in Cacodylate buffer 0.05M (pH7.4), 0.09M sucrose, and 0.9mM MgCl2. Briefly, specimens were postfixed in 1% osmium tetroxide, 0.8% potassium ferrocyanide in cacodylate buffer (0.1 M, pH 7.2), treated with 0.2% Tannic Acid in 0.05M cocadylate buffer (pH 7.0), stained with 1% uranyl acetate in Sodium Acetate (pH 5.2), dehydrated in an alcohol series, and embedded in Hard EPON. 50-nm sections were imaged at 23,000x magnification using a TecnaiSpirit (FEI) operated at 80 kV and equipped with an Eagle CCD camera (FEI). Using the relative position of GFP::SAS-7 foci and neighboring nuclei in fluorescence images facilitated the search for centrioles in EM serial sections. Gray values were adjusted and Gaussian blur filtering 1.5 was applied on the displayed EM images. The two gonads analyzed were straightened using the “Straighten” Fiji plugin (KOCSIS et al., 1991) and the distance of each nucleus studied to the syncytium exit was determined. For measurements of ultrastructural elements, each data point is the average of 4 measurements extracted from lines drawn along the height of the feature. In some cases, ultrastructural elements could not be measured because they were not visualized accurately, the view of the centriole was too tilted or it was not apparent anymore due to ongoing elimination.

### Quantification and statistical analysis

For Fig.1 and Fig.2, correlation between the 2 markers for each condition was determined taking into account the 12 or 15 positions respectively. R^2^ of fits are mentioned in the legends. Levene’s test for equality or inequality of variance was performed by using: https://www.socscistatistics.com/tests/levene/default.aspx. Required normality of input data was tested for and confirmed to be not significantly different from normal distribution by performing a Kolmogorov-Smirnov test (p-value for WT: 0.797 and *sas-1(ts)*: 0.993) using: https://www.socscistatistics.com/tests/kolmogorov/default.aspx

**Movie S1. Centriolar focus movement from the nuclear envelope to the plasma membrane** Widefield time lapse imaging of an oocyte from a worm immobilized in the microfluidic device and expressing MEL-28::GFP as well as RFP::SAS-7. Top: brightfield; bottom: max projection of relevant optical slides of the merge between MEL-28::GFP (green) and RFP::SAS-7 (magenta). Note that the RFP::SAS-7 centriolar focus detaches from the nuclear envelope at 0 min, then migrates to the plasma membrane, which is reached at 52 min, before the foci becoming undetectable at 166 min.

