## supplemental data for "Centriole elimination during *C. elegans* oogenesis initiates with loss of the central tube protein SAS-1"

**SUPPLEMENTARY FIGURES LEGENDS**

**Figure S1. Centriolar and PCM components are lost progressively starting in late meiotic prophase.**

**A-D.** Gonads expressing RFP::SAS-7 (magenta) and the indicated other protein tagged with GFP (green). In inset for -9 oocyte in C, the strong green signal corresponds to plasma membrane GIP-1.

**E, F.** Gonads expressing AIR-1::GFP (E) and ZYG-9::GFP (F) together with RFP::SAS-7. Numbered white rectangles mark magnified insets (right). Arrowheads point to SAS-7 foci (magenta) and their corresponding position in the green channel. AIR-1 and ZYG-9 enrichment at centrioles is restricted to mitosis.

**Figure S2. Movement of centriolar protein foci from the nucleus to the plasma membrane is actin- and sperm-independent, but partially kinesin-dependent.**

**A-D.** Gonads from control (F) and *fem-1(hc17ts)* (H) worms expressing RME-2::GFP and RFP::SAS-7, together with corresponding quantifications (G, I). In the insets, dashed circles mark nuclei position and arrowheads point to RFP::SAS-7 foci. Note that centriolar focus detachment and movement occurs in *fem-1(hc17ts)* despite the absence of sperm in this mutant. N: G=11, I=12

**E-I.** Quantifications in the indicated RNAi treatments of the percentage of oocytes bearing a SAS-7 focus (SAS-7 total, grey lines) among which those with RFP::SAS-7 detached foci (SAS-7 detached, magenta dashed lines) as well as of oocytes with RME-2::GFP enriched at the plasma membrane (green dashed lines), relative to their position in the gonad. E-G, as well as H and I, are from independent experiments. N: A=12; B=18; C=17, D=9; E=18.

**Figure S3. Depletion of PLKs or PCM does not lead to precocious elimination in *C. elegans***

**A, B.** Gonads from worms of indicated genotype expressing SPD-5::GFP, RFP::SPD-2 and RME-2::GFP. The double mutant gonad is highly disorganized, yet RME-2::GFP is enriched at the plasma membrane of cells close to the spermatheca (-1 to -4); moreover, RFP::SPD-2 and SPD-5::GFP foci are present. N: A=13; B=12.

**C, D.** Immunostaining of extruded gonads from worms of indicated genotypes; centrioles (IFA-1 antibodies): red; DNA: cyan. Asterisks mark somatic sheath cell nuclei. Brackets indicate pairs of centriolar foci in the vicinity of the same nucleus.

**E, F.** Immunostaining of dissected gonads from control and *pcmd-1(t3421ts)* worms at the restrictive temperature expressing RFP::SPD-5 and GFP::SAS-7 stained with antibodies against GFP (cyan) and SAS-4 (yellow); DNA: grey. RFP fluorescence (magenta) persists after fixation, which allow to visualize SPD-5 in the control condition. Note loss of cytoplasmic signal in -3 and -1 oocytes, which sometimes happen upon fixation. In F, asterisk marks unspecific SAS-7 staining.

**Figure S4. SPD-5 and α-tubulin intensities change concomitantly during prophase progression.**

**A.** U-Ex-STED images of centrioles top-views over the course of prophase stained for α-tubulin and SPD-5. The LUT "Fire" represent changes of fluorescence intensity over prophase progression (from bright to dim: white, yellow, red, purple, blue and black).

**B.** Changes of normalized centriolar SPD-5 fluorescence intensity relative to α-tubulin during prophase progression imaged by U-Ex-STED microscopy. All values were background subtracted and first normalized to the α-tubulin signal intensity in the same centriole and then normalized to the mean of early prophase measurements of each series. N (early prophase, late pachytene and diplotene): 18, 30 and 18. The middle lines of the box-plots correspond to the median, the cross represents the mean, the box includes 50% of values (IQR) and the whiskers show the range of values within 1.5*IQR.

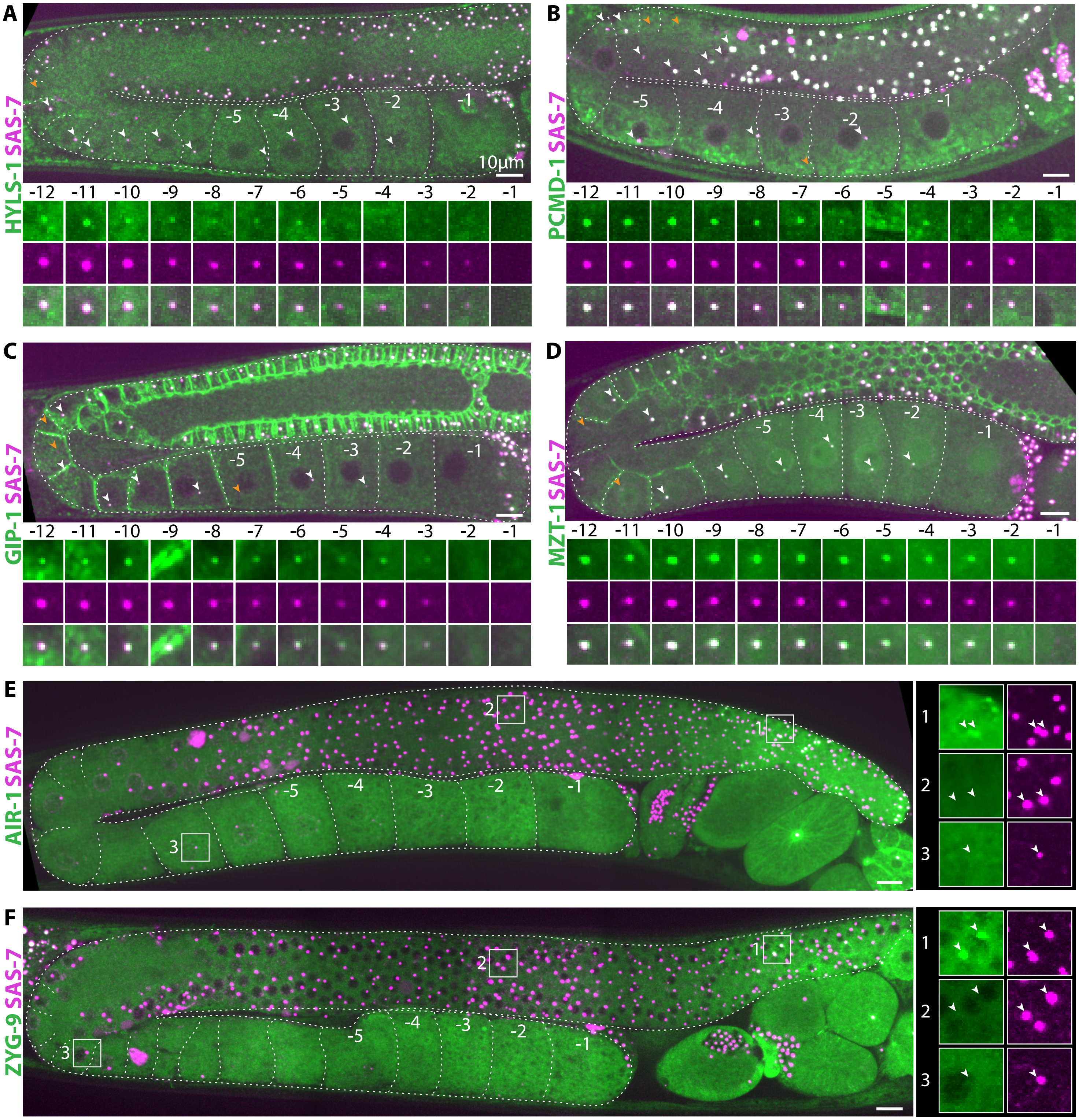
A
HYLS-1 SAS-7
10µm
-12 -11 -10 -9 -8 -7 -6 -5 -4 -3 -2 -1
B
PCMD-1 SAS-7
C
GIP-1 SAS-7
D
MZT-1 SAS-7
E
AIR-1 SAS-7
F
ZYG-9 SAS-7
1
2
3

FIGURE S1

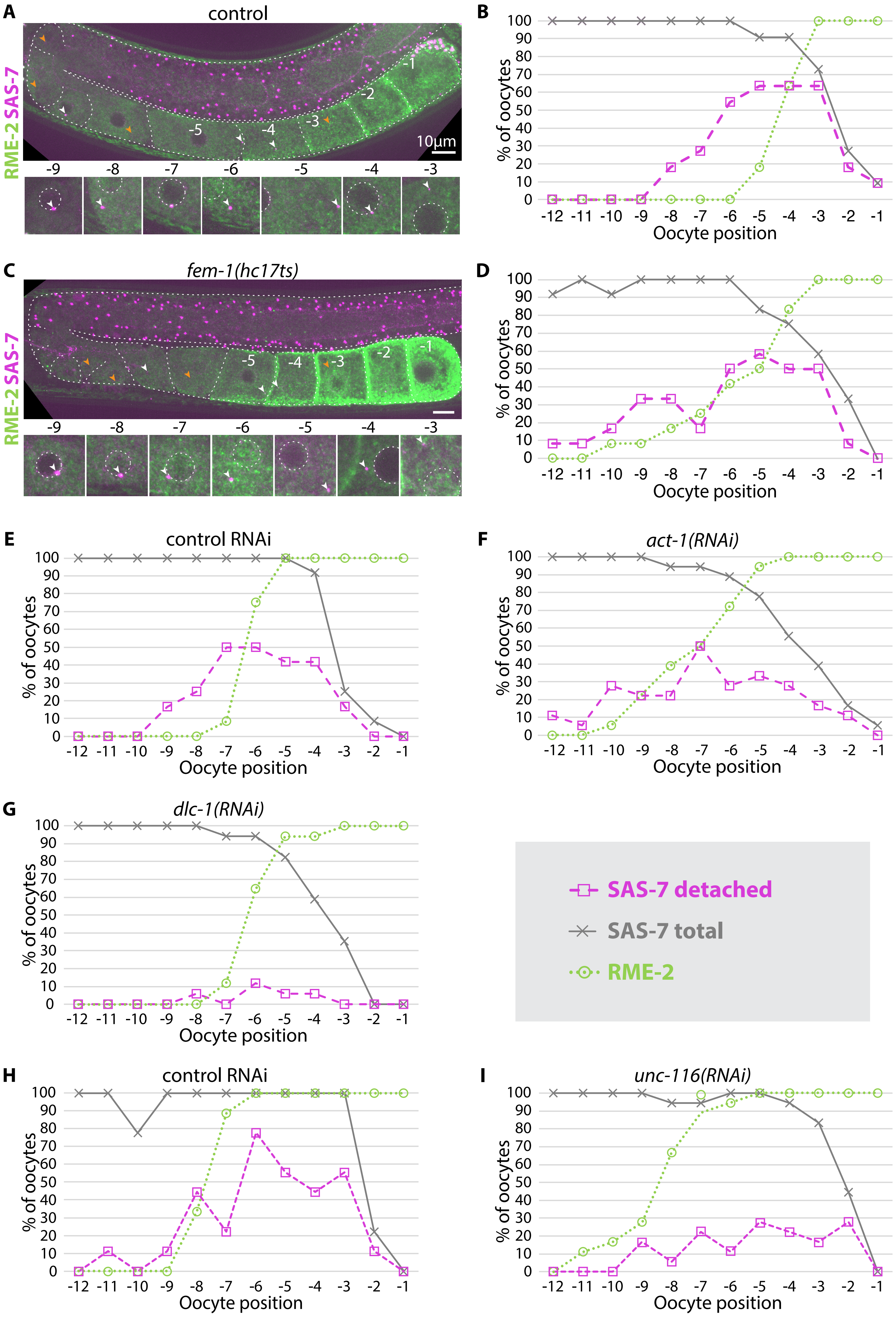
A
control
RME-2 SAS-7
10µm
-9 -8 -7 -6 -5 -4 -3
B
% of oocytes
Oocyte position
C
*fem-1(hc17ts)*
RME-2 SAS-7
D
% of oocytes
Oocyte position
E
control RNAi
% of oocytes
Oocyte position
F
*act-1(RNAi)*
% of oocytes
Oocyte position
G
*dlc-1(RNAi)*
% of oocytes
Oocyte position
SAS-7 detached
SAS-7 total
RME-2
H
control RNAi
% of oocytes
Oocyte position
I
*unc-116(RNAi)*
% of oocytes
Oocyte position

**FIGURE S2**

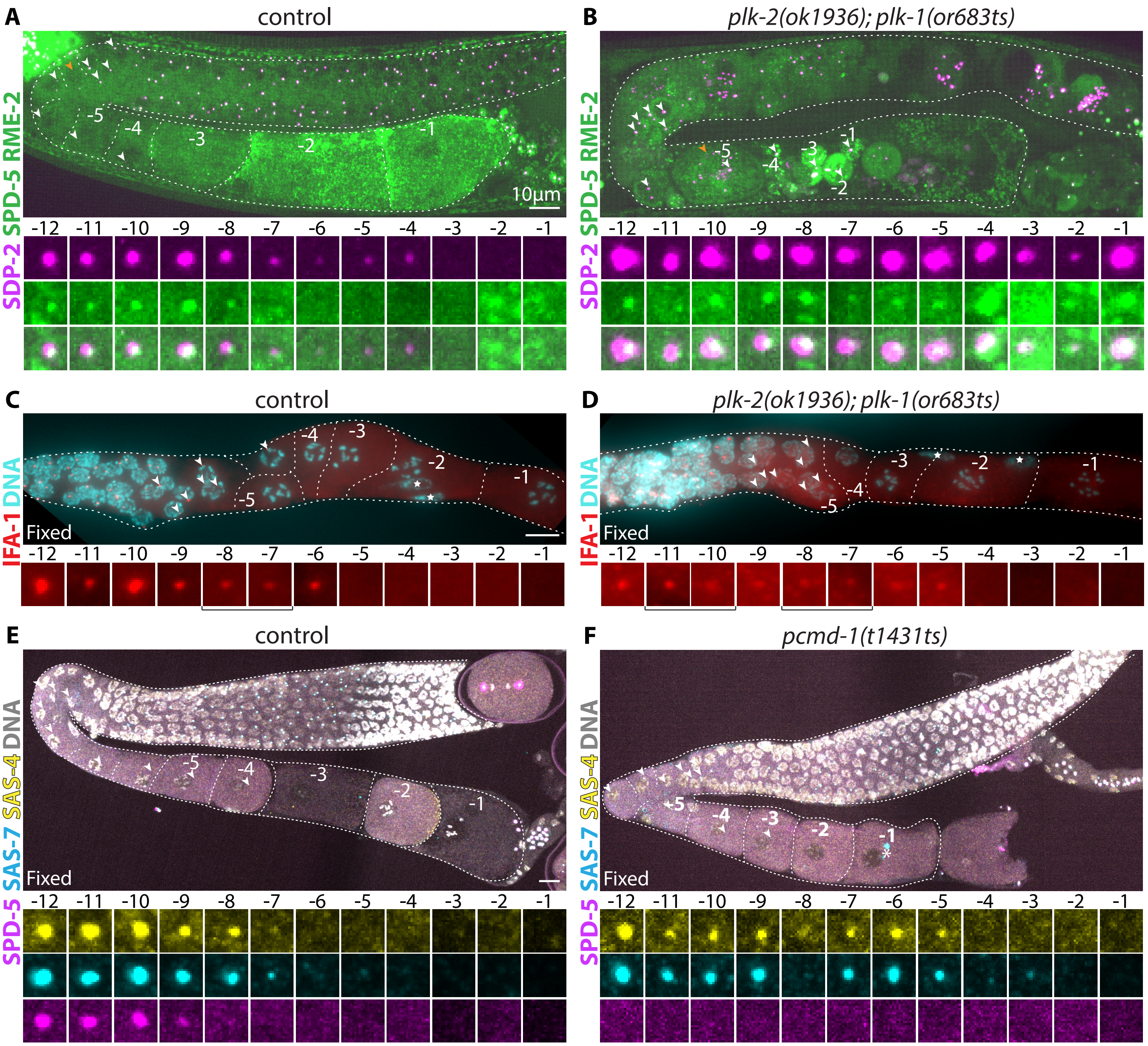

A
control
B
plk-2(ok1936); plk-1(or683ts)
SPD-5 RME-2
SDP-2
10µm
-12 -11 -10 -9 -8 -7 -6 -5 -4 -3 -2 -1
C
control
D
plk-2(ok1936); plk-1(or683ts)
IFA-1 DNA
Fixed
E
control
F
pcmd-1(t1431ts)
SPD-5 SAS-7 SAS-4 DNA
Fixed


**Figure S3**

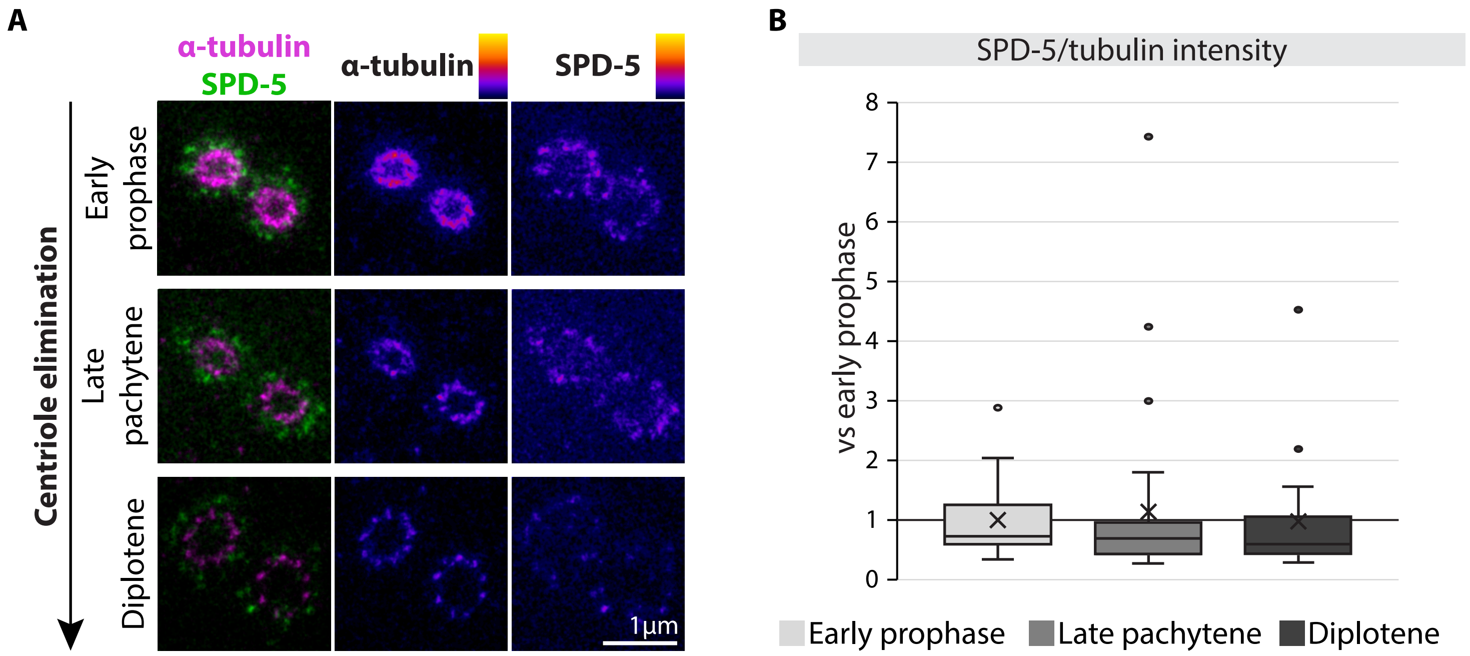
A
α-tubulin
SPD-5
α-tubulin
SPD-5
Centriole elimination
Early prophase
Late pachytene
Diplotene
1µm
B
SPD-5/tubulin intensity
vs early prophase
8
7
6
5
4
3
2
1
0
Early prophase
Late pachytene
Diplotene

**FIGURE S4**
